# Quantitative cell morphology in *C. elegans* embryos reveals regulations of cell volume asymmetry

**DOI:** 10.1101/2023.11.20.567849

**Authors:** Guoye Guan, Zelin Li, Yiming Ma, Jianfeng Cao, Ming-Kin Wong, Lu-Yan Chan, Hong Yan, Chao Tang, Zhongying Zhao

## Abstract

The dynamics of cellular morphology throughout development are crucial for morphogenesis and organogenesis, yet their systematic characterization remains a significant challenge. By integrating both nuclear position and advanced cell membrane labeling, we develop a novel method that enables the segmentation of surfaces for over 95% of cells produced during *Caenorhabditis elegans* embryogenesis. With this method, we segment eight wild-type and four perturbed embryos. The output, including cell identity, shape, volume, surface and contact area, can be visualized using our custom software. We demonstrate that signaling interactions, such as Notch and Wnt, regulate not only the asymmetry of cell fate but also the asymmetry of cell volume in conjunction with mechanical compression. Furthermore, we find that the asymmetries of fate and volume are generally interconnected.

**ONE SENTENCE SUMMARY:** **Systematic quantification of cell morphology with resolved cell lineage and cell fate in developing *C. elegans* embryo uncovers multimodal regulations of cell size.**

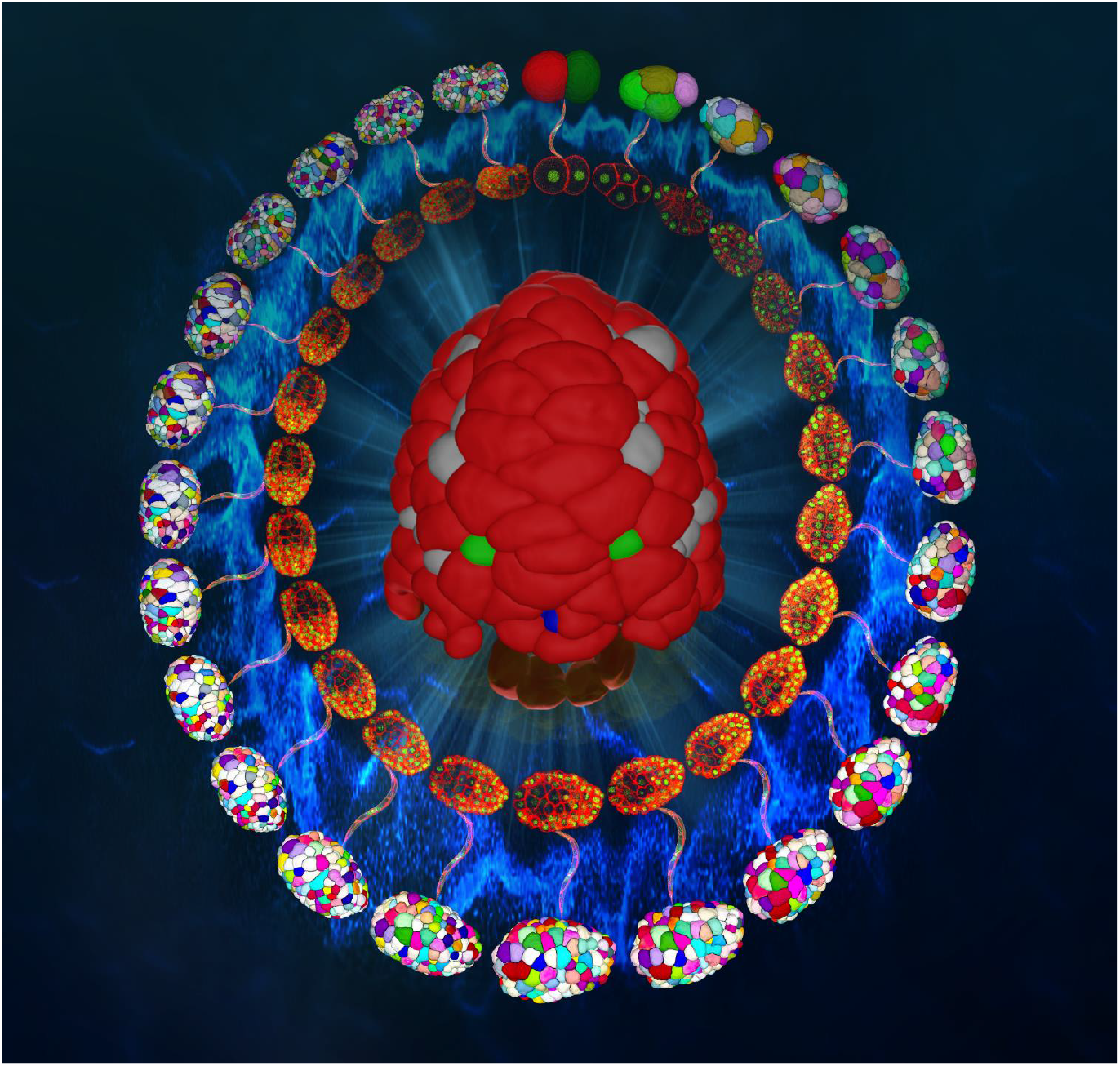

**Potential Cover/Featured Image: Smiling “ghost” face of *C. elegans* embryo.** Center: shown are skins (colored in red, dorsal view with anterior to the bottom) of a 400-celled *C. elegans* embryo. Cells not covered by the skins are colored in gray, green, blue, and transparent yellow, respectively. Internal and external circles: confocal fluorescence images (GFP for cell nucleus and mCherry for cell membrane) and reconstructed cell morphologies of the *C. elegans* embryo from the 2- to 550-cell stages respectively, connected with images of a *C. elegans* adult expressing the same markers.

## INTRODUCTION

Cell morphology, usually defined as its spatial boundary composed of the plasma membrane (hereafter referred to as cell membrane), consists of significant information including but not limited to cell shape, volume, surface and contact area [Cao *et al., Nat. Commun.*, 2020]. Among them, the shape of a cell is known as an indicator of its physiological states like fate and function [Luxenburg *et al., Exp. Cell Res.*, 2019; Eddy *et al., Sci. Rep.*, 2021], while multiple cells with specific shapes can collectively constitute the more complex and functional morphology at the tissue, organ, and individual scales [Gómez -Gálvez *et al., Nat. Commun.*, 2018; Witvliet *et al., Nature*, 2021; Guignard *et al., Science*, 2020]. Besides, the cell volume, surface and contact area are also critical for numerous biological events. For example, the heterogeneous cell volume could be significantly correlated with heterogeneous cell cycle length, coordinating the movements, positions, and contacts of cells in concert [Fickentscher *et al., Phys. Rev. Lett.*, 2016]; then, some ligands and receptors accumulated on the surface of specific cells can bind to each other through cell-cell contact for signaling transduction and fate specification [Priess, *WormBook*, 2005]. For the reasons given above, the cell morphology reconstruction in different organisms, especially for the multicellular and developing ones, has been a popular focus not only in computer science, but also in cell and developmental biology, and so forth [Guan *et al., Comput. Struct. Biotechnol. J.*, 2022].

Nowadays, the cell morphology in the embryogenesis of many species (*e.g.*, nematode, ascidian, fruitfly, zebrafish) has been a reconstruction target, as it characterizes lots of landmark morphogenesis and organogenesis where a number of cells are migrating, dividing, and differentiating simultaneously [Azuma *et al., BMC Bioinform.*, 2017; Guignard *et al., Science*, 2020; Stegmaier *et al., Dev. Cell*, 2016]. Among them, the nematode *Caenorhabditis elegans* (*abbr., C. elegans*) is a widely used model animal for various research themes because of its highly invariant developmental programs at the cellular level, with respect to space, time, lineage, and fate [Sulston *et al., Dev. Biol.*, 1983]. Compared to the other organisms, its embryogenesis ends with a constant cell number (*i.e.*, 558 living cells in hermaphrodite prior to hatching) and condensed cell fate assignments; for instance, there are only 20 cells forming its gut. Thus, researchers in all fields can take advantage of the behavior of a few particular cells for studying critical biological phenomena, like gastrulation and apoptosis [Nance *et al., Development*, 2002; Cordes *et al., Development*, 2006]. In recent years, both experimental and computational approaches customized for the *C. elegans* embryo have been kept ungraded for accurately reconstructing its complete morphological map over development and at the single-cell resolution [Azuma *et al., BMC Bioinform.*, 2017; Chen *et al., Genetics*, 2018; Cao *et al., BMC Bioinform.*, 2019; Xiong *et al., PLoS ONE*, 2020; Cao *et al., Nat. Commun.*, 2020; Thiels *et al., Bioinformatics*, 2021].

Till now, a lot of attention and effort has been paid to the cell morphology reconstruction for *C. elegans* embryogenesis, where two major aspects are always the key to good performance: continuously distinguishable fluorescence signal on the cell membrane and the segmentation algorithm that is able to unambiguously recognize it. However, due to the rapid cleavage along with drastically decreasing cell size, the volumetric pixels with a constant resolution for characterizing the membrane or morphology of a cell are less and less, so the fluorescence signal becomes blurred gradually [Cao *et al., Nat. Commun.*, 2020]. To this day, despite that the cell morphology at the whole-embryo level can be reconstructed for the first half of embryogenesis (*i.e.*, roughly from the 1-to 350-cell stages), the second half (*i.e.*, roughly from the 350-to 550-cell stages) cannot be accurately reconstructed no matter which previously-proposed *C. elegans*-customized algorithm is employed [Azuma *et al., BMC Bioinform.*, 2017; Cao *et al., BMC Bioinform.*, 2019; Cao *et al., Nat. Commun.*, 2020; Thiels *et al., Bioinformatics*, 2021]. Since the late stages cover a lot of developmental landmarks (*e.g.*, ventral cleft closure, dorsal intercalation, epidermal enclosure) [Chisholm *et al., WormBook*, 2005], its cell-resolved morphological map is urgently needed by researchers in all kinds of fields.

In this paper, we established a new platform that allows qualitative and quantitative analysis of cell shape, volume, surface and contact area in *C. elegans* embryos at up to the 550-cell stage, when most embryonic cells complete their last round of division with terminal fate. The platform performs significantly better than the state-of-the-art methods in segmenting cell membranes beyond the 350-cell stage of *C. elegans* embryo. We demonstrated that cell fate asymmetry is commonly coupled with cell volume asymmetry, and cell signaling such as Notch signaling not only regulates the fate asymmetry, but also controls the volume asymmetry of the relevant cells, and successive Notch signaling events contribute to the remarkably large volume of the kidney cell in *C. elegans* embryos.

### Automated cell segmentation aided by the fluorescence labeling of cell nucleus and cell membrane

Reconstructing embryonic cell morphology is a time-consuming and resource-intensive task, which requires a methodology to segment all the cell membranes systematically and automatically. To comprehensively reconstruct the *C. elegans* embryonic morphological map, especially for the late developmental stages (*e.g.*, the 550-cell stage), in this work, we present an automatical pipeline, *CMap*, to construct digitized *C. elegans* embryos with high segmentation accuracy and high computational speed (Table S1), by reconstructing all embryonic cell morphologies at each time point, building the whole map of cell lineages and cell fates for every embryo, and extracting statistical morphological features for each cell (Figure 1A; Movie S1; Materials and Methods - Three-dimensional illustration for segmented cell morphologies). We first construct a novel strain with more distinct and homogeneous fluorescence labeling with mCherry on the cell membrane at both embryonic and postembryonic stages (Figure 1B; Figure S1; Materials and Methods - Worm strains and maintenance; Materials and Methods - Fluorescence microscopy for embryonic stages; Materials and Methods - Fluorescence microscopy for postembryonic stages). Meanwhile, the fluorescence labeling with GFP (Green Fluorescent Protein) on cell nucleus in another channel is tracked using the software *StarryNite* and *AceTree*, providing the cell nuclear positions as well as cell identities, cell lineages, and cell fates, with the aid of previous knowledge (Materials and Methods - Automated cell nucleus tracing and lineaging) [Murray *et al., Nat. Protoc.*, 2006; Sulston *et al., Dev. Biol.*, 1983; Li *et al., Cell Rep.*, 2019]. The cell nuclear positions are subsequently used as alternative seeds for the single cell morphology reconstruction by an advanced adaptive deep convolutional neural network called EDT-DMFNet (Euclidean Distance Transform Dilated Multi-Fiber Network) for cell membrane recognition (Figure S2), which successfully handles the late developmental stages with the relatively larger cell number and smaller cell size (Figure 1B). This single-cell segmentation serves for accurate downstream morphological analyses, where the morphological features of the individual and lineage- and fate-wise 3D cell objects are automatically analyzed across the whole predicted embryonic morphology (Movie S2; Movie S3; Materials and Methods - Calculation of cell volume, cell surface area, and cell-cell contact area). Notably, with an NVIDIA 2080 Ti GPU (Graphics Processing Unit), *CMap* takes less than 3 hours on average to implement automatic cell segmentation for a *C. elegans* embryo from the 4-to 550-cell stages (Table S1), allowing high-throughput processing for both normal and perturbed (*e.g.*, with RNA interference, mechanical compression, eggshell removal, laser ablation) embryos in the future [Ho *et al., Mol. Syst. Biol.*, 2015; Jelier *et al., Cell Syst.*, 2016; Fickentscher *et al., Sci. Rep.*, 2017; Chen *et al., Genetics*, 2018].

**Fig. 1.**
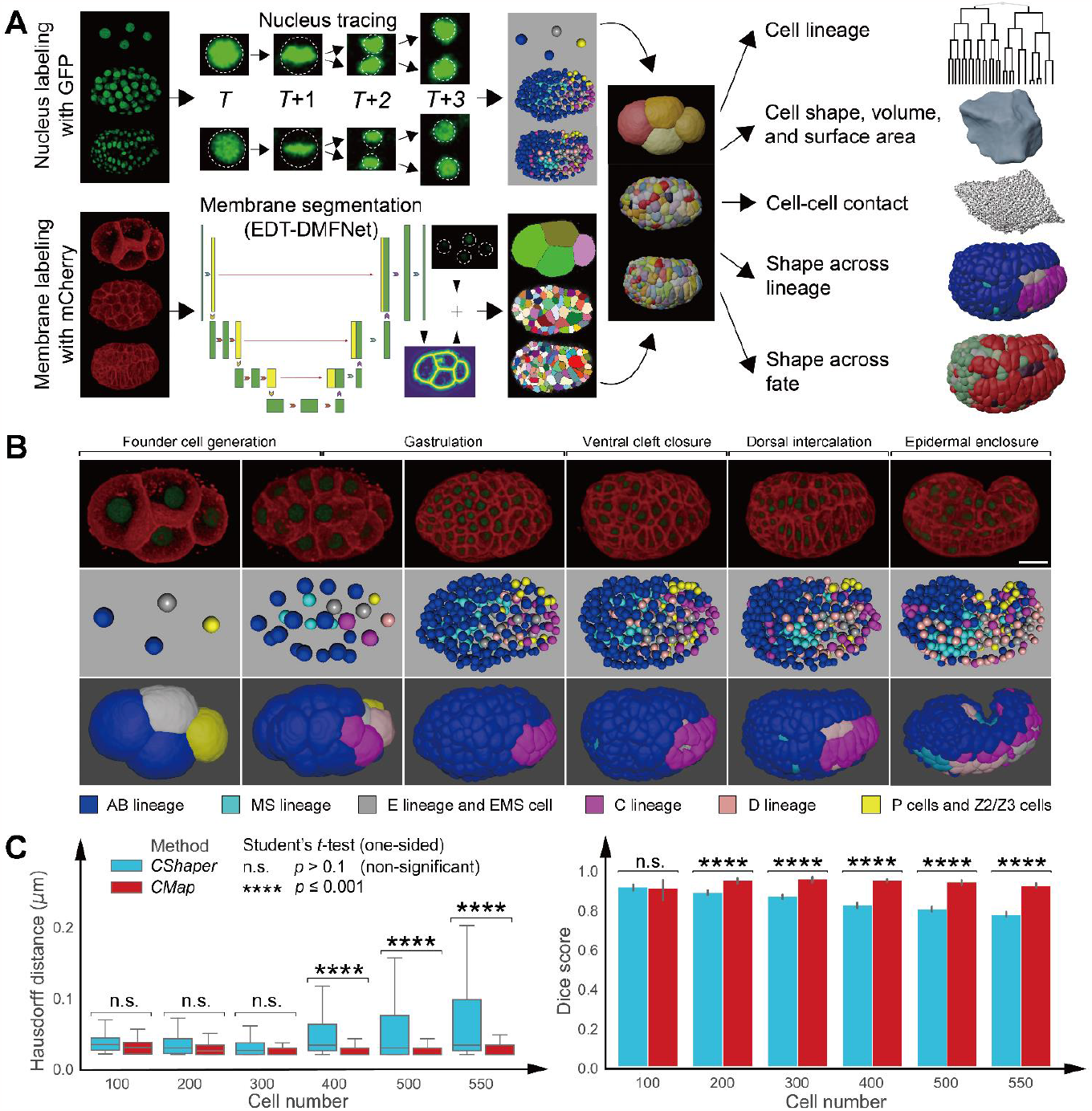
The cell morphology reconstruction for *C. elegans* embryogenesis by *CMap*. (**A**) Data processing pipeline of *CMap*, with the input of fluorescence labeling cell nucleus and cell membrane and output of cell-resolved morphological map including individual and lineage- and fate-wise cell shape, volume, surface and contact area. (**B**) Fluorescence imaging (upper row; with a scale bar of 10 μm plotted in the bottom right corner), cell nucleus tracing (middle row), and cell membrane segmentation (lower row) throughout *C. elegans* embryogenesis from the 4- to >550-cell stages, highlighted with the developmental landmarks marked by their corresponding typical cell number documented in [Chisholm *et al., WormBook*, 2005]. (**C**) Performance evaluation for *CShaper* and *CMap* by Hausdorff distance (left panel) and Dice score (right panel) at different developmental stages (represented by cell numbers), using 12 manually annotated 3D images as the benchmark. Here, for the Hausdorff distance, the lower and upper quartiles of a data group are shown with a colored box, with its median labeled inside and two whiskers extended to the maximum and minimum data value; for the Dice score, the average of a data group is shown by a colored column, with all the data values plotted with gray dots; the statistical significance is obtained by the one-sided Student’s *t*-test and is listed at the top.

### Outperformance of *CMap* compared to other state-of-the-art algorithms

The convolutional-neural-network-based method proposed in this work, *CMap* (Figure 1A), consists of two algorithmic advances for processing the denoised 3D fluorescence images: 1. the deep learning network with optimal loss function and network structure (Materials and Methods - Automatic segmentation; Materials and Methods - Training data augmentation); 2. long-term manually-assisted cell nucleus tracking that provides seeds for cell membrane recognition (Materials and Methods - Automated cell nucleus tracing and lineaging). To evaluate how close the cell morphology reconstructed by *CMap* is to the real one, we manually segment the 3D images with fluorescently-labeled cell membranes at six different time points (at around 100-, 200-, 300-, 400-, 500-, and 550-cell stages) in two embryo samples (Materials and Methods - Manual annotation). Using those ground truths (4046 cell objects in total) as a reference, we compare the performance between *CMap* and five state-of-the-art cell segmentation algorithms — *CShaper* [Cao *et al., Nat. Commun.*, 2020], *3DCellSeg* [Wang *et al., Sci. Rep.*, 2022], *CellPose3D* [Eschweiler *et al., ICIP*, 2022], *StarDist3D* [Weigert *et al., WACV*, 2020], and *VNet* [Milletari *et al., 3DV*, 2016], in order to study the advantages and disadvantages (Materials and Methods - Performance evaluation of *CMap* and other existing cell segmentation algorithms).

First, *CShaper* is separately compared to *CMap*. In our paper published three years ago [Cao *et al., Nat. Commun.*, 2020], *CShaper* was devised for segmenting the living *C. elegans* embryo for its first half embryogenesis (*i.e.*, roughly from the 1-to 350-cell stages) in the mechanically compressed state (for a narrow light field), with fluorescence labeling on cell membrane and watershed algorithm based on automatic seeding without cell nuclear position involved; its performance in cell morphology reconstruction drops sharply after the 350-cell stage, as the cell number keeps increasing and the cell size keeps decreasing over cleavage [Guan *et al., Comput. Struct. Biotechnol. J.*, 2022]. Here, *CShaper* is trained under the same conditions as the ones for *CMap* (Materials and Methods - Automatic segmentation; Materials and Methods - Training data augmentation; Materials and Methods - Manual annotation). Intuitively shown by the segmented cell morphologies, when the embryo is imaged without mechanical compression and with its natural eggshell shape, *CShaper* fails to deal with the embryo periphery as soon as at ∼100-cell stage, and almost each cell acquires an unsmooth shape at the 550-cell stage, compared to both the ground truth and *CMap* outputs (Figure S3; Figure S4); this may be attributed to the wider light field for embryo imaging in the mechanically uncompressed state, which amplifies the attenuation and scattering of the laser. Quantitatively evaluated by the Hausdorff distance (defined as the largest of all distances from a pixel in one region to the closest pixel in the other region) and Dice score (defined as the ratio between the overlapping region and the overall region) used before [Cao *et al., Nat. Commun.*, 2020], on average, *CMap* achieves a Hausdorff distance of 0.0241 ± 0.0043 μm smaller than the one of *CShaper* (0.0518 ± 0.0098 μm) and a Dice score of 0.9426 ± 0.0242 larger than the one of *CShaper* (0.8510 ± 0.0519). The significant superiority regarding both criteria (one-sided Student’s *t*-test, *p* ≤ 0.001) holds since the 400-cell stage, suggesting that *CMap* indeed has better cell segmentation performance than *CShaper*, especially for the second half of *C. elegans* embryogenesis (Figure 1C).

Second, among the other four general cell segmentation algorithms (*i.e., 3DCellSeg, CellPose3D, StarDist3D, VNet*), *StarDist3D* and *VNet* have a Hausdorff distance approaching the ones of *CShaper* and *CMap* at up to the 300- and 200-cell stage respectively, revealing their applicability for the first half of *C. elegans* embryogenesis even though it’s not customized for this system (one-sided Student’s *t*-test, *p* > 0.1). Nonetheless, *CMap* still exhibits significantly better performance than all the other considered algorithms at the 400-, 500-, and 550-cell stages, no matter for Hausdorff distance or Dice score (one-sided Student’s *t*-test, *p* ≤ 0.05) (Figure S5; Table S2).

### Quantitative morphological map for nearly all cells produced during *C. elegans* embryogenesis

To establish a cell-resolved morphological map for *C. elegans* embryogenesis with both statistical reliability and data completeness, a total of eight *C. elegans* wild-type embryos (marked as “WT_Sample1” ∼ “WT_Sample8”) with GFP labeling their cell nuclei and mCherry labeling their cell membranes are imaged for at least four hours at an interval of ∼1.5 minutes (Table S3; Materials and Methods - Worm strains and maintenance; Materials and Methods - Fluorescence microscopy for embryonic stages), followed by cell lineaging based on the software *StarryNite* and *AceTree* (Materials and Methods - Automated cell nucleus tracing and lineaging; Materials and Methods - Quality control on embryo imaging and cell tracking) [Murray *et al., Nat. Protoc.*, 2006]. For every embryo sample, the cell tracking starts no later than the 4-cell stage and ends after the 550-cell stage, when all the 48 final progenies in the C lineage have existed for at least five time points (Figure 2A; Figure 2B) [Sulston, et al. *Dev. Biol.*, 1983]; eventually, over 95% of cells in *C. elegans* embryogenesis have appeared and been recorded in at least one embryo sample (Table S4) [Sulston *et al., Cold Spring Harbor Laboratory Press*, 1988]. Within these embryo samples, there are 1292 unidentical cells detected in total, including 1188 non-apoptotic cells and 104 cells with programmed death (92.04% of all the 113 *C. elegans* hermaphrodite embryonic apoptotic cells). Among them, 1190 unidentical cells are recorded in every embryo, of which 589 have a complete lifespan (*i.e.*, cell cycle) and 79 have programmed cell death. Compared to our previously published dataset generated by *CShaper*, which focused on the 4-to 350-cell stages, the content of the current dataset generated by *CMap* is almost double and provides sufficient information on the second half of *C. elegans* embryogenesis (*i.e.*, roughly from the 350-to 550-cell stages) (Table S5). For the 3D cell regions or data points (*c, j, T*) marked by cell identity *c*, embryo sample *j*, and time point *T*, an amount of 395741 cell regions (99.99%) are successfully segmented by *CMap*, including 5905 cell regions with two cell nuclei inside, *i.e.*, proceeding cytokinesis (Table S4). Subsequently, 10.89% of the cell regions are identified with abnormal volume or shape (Figure S6; Figure S7; Figure S8; Materials and Methods - Data filtering based on abnormal cell volume and cell shape). For those aberrant cell regions, 69.10% are non-apoptotic cells and presumably have a nearly unchanged volume over time as well as a continuous space occupation, so they are removed; the remaining ones (13320 in total) are the apoptotic cells which possibly have a sudden volume decrease, so they are manually checked and corrected by ten technicians (Figure S9; Figure S10) [Driscoll *et al., Proc. Natl. Acad. Sci. U. S. A.*, 2017]. At last, a *C. elegans* embryonic morphological map with a total data point loss rate lower than 8% is established with confident data quality, faithfully providing a highly reproducible and statistical source to study all kinds of cell morphology-related problems (Table S6; Table S7; Table S8).

**Fig. 2.**
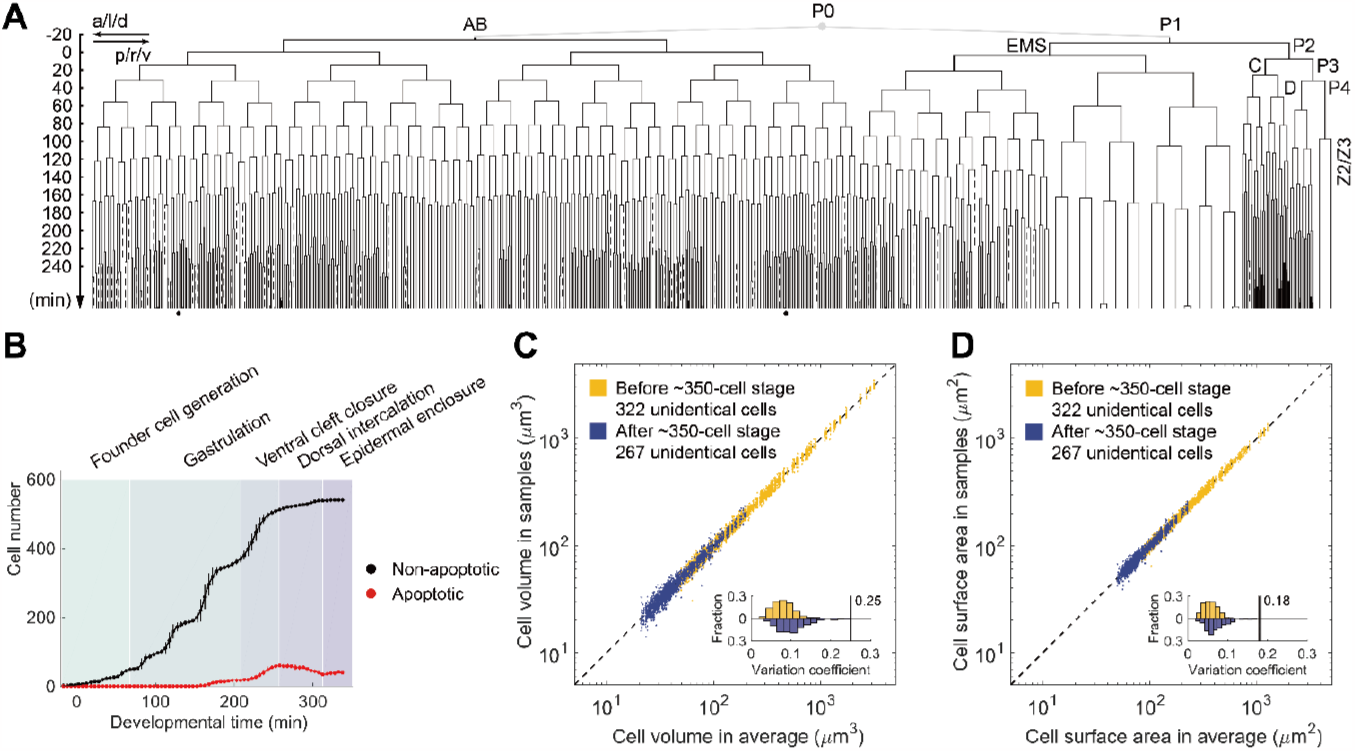
The quantitative data in the morphological map covering the majority of *C. elegans* embryogenesis. (**A**) The embryonic cell lineage tree averaged over the eight *C. elegans* wild-type embryo samples. Here, the solid and dashed lines represent non-apoptotic and apoptotic cells respectively, while the cells failing in segmentation in all embryo samples are denoted by the dots beneath. (**B**) The cell number (with the solid circle representing the average and the vertical line representing the standard deviation within a constant interval) over developmental time, where the duration between developmental landmarks marked by their corresponding typical cell number documented in [Chisholm *et al., WormBook*, 2005] are illustrated using different colors in the background. (**C**) The consistency between average cell volume (horizontal coordinate) and its values in the eight *C. elegans* wild-type embryo samples (vertical coordinate). Here, each dot denotes the average volume (horizontal coordinate) over eight embryo samples and the value in one embryo sample for a unique cell; the 322 unidentical cells before the ∼350-cell stage and the 267 unidentical cells after the ∼350-cell stage are represented by blue and yellow respectively, while the distributions of their correlation coefficients are shown in the inset. (**D**) The consistency between average cell surface area (horizontal coordinate) and its values in the eight *C. elegans* wild-type embryo samples (vertical coordinate). Here, each dot denotes the average volume (horizontal coordinate) over eight embryo samples and the value in one embryo sample for a unique cell; the 322 unidentical cells before the ∼350-cell stage and the 267 unidentical cells after the ∼350-cell stage are represented by blue and yellow respectively, while the distributions of their correlation coefficients are shown in the inset.

The straightforward morphological feature outputted by *CMap* cell segmentation is the shape of a cell; Figure S11 displays the time-lapse shape change of representative cells from different lineages and with different fates since the ≥550-cell stage as well as the germline cells. Apart, three more quantitative morphological features are outputted: cell volume (*i.e., V*), cell surface area (*i.e., A*_S_), and cell-cell contact area (*i.e., A_C_*) (Figure 1A; Materials and Methods - Calculation of cell volume, cell surface area, and cell-cell contact area). Among them, the control on cell size (*incl.*, cell volume and cell surface area) is of high importance in the classic concept of how accurate *C. elegans* embryogenesis is, and many studies have been carried out to measure its accuracy as well as to figure out its downstream regulatory targets in both biological and physical contexts [Cao *et al., Nat. Commun.*, 2020; Jankele *et al., eLife*, 2021; Fickentscher *et al., Phys. Rev. Lett.*, 2016]. For every *C. elegans* wild-type embryo sample in our dataset, in addition to the 322 unidentical cells with a complete lifespan recorded before the ∼350-cell stage [Cao *et al., Nat. Commun.*, 2020], there are 289 more unidentical cells recorded with a complete lifespan after that; it’s found that both the early and late cells exhibit considerable size (*incl.*, cell volume and cell surface area) control among embryo samples, with a correlation coefficient always smaller than 0.25, while the late cells show higher relative fluctuation (Figure 2C; Figure 2D).

### Lineage- and fate-wise characterization of morphogenesis and organogenesis at the cellular resolution

Based on the invariant cell lineage programs and cell fate assignments in *C. elegans* embryogenesis [Sulston *et al., Dev. Biol.*, 1983], cell shapes across tissue-, organ-, and embryo-scales with cell fate (*incl.*, neuron, pharynx, intestine, germline, muscle, skin, death, unspecified, others) labeled is generated (Figure 3A; Movie S2; Movie S3). It should be pointed out that, starting from the cell fate categorization summarized in [Li *et al., Cell Rep.*, 2019], if the two daughters of a cell have different cell fates, it’ll be categorized into “unspecified”; if the two daughters of a cell have the same cell fate, it’ll be categorized into its daughters’ cell fate; if the fate of a cell has been determined, its two daughters will be categorized into its cell fate (Table S4). After that, the stereotypic anatomical structure of different tissues and organs inside an embryo can be observed intuitively from different views (Movie S4); such observation is achieved with a data point loss rate lower than 5% for all the tissues and organs at all the ∼100-, ∼200-, ∼300-, ∼400-, ∼500-, and >550-cell stages, as shown in Figure S12. Taking the mesoderm and endoderm as representatives: the neuron and muscle (mesoderm) are distributed symmetrically on the left and right sides of the embryo (shown in the 3^rd^ and 5^th^ columns of Figure 3A); the pharynx and intestine (endoderm) form a tunnel through the anterior and posterior of the embryo, where the intestine is skewed around the germline cells Z2 and Z3 with roughly 90°, probably serving as protection for them in consideration of their essential role in reproduction (shown in the 4^th^ column of Figure 3A; Movie S5).

**Fig. 3.**
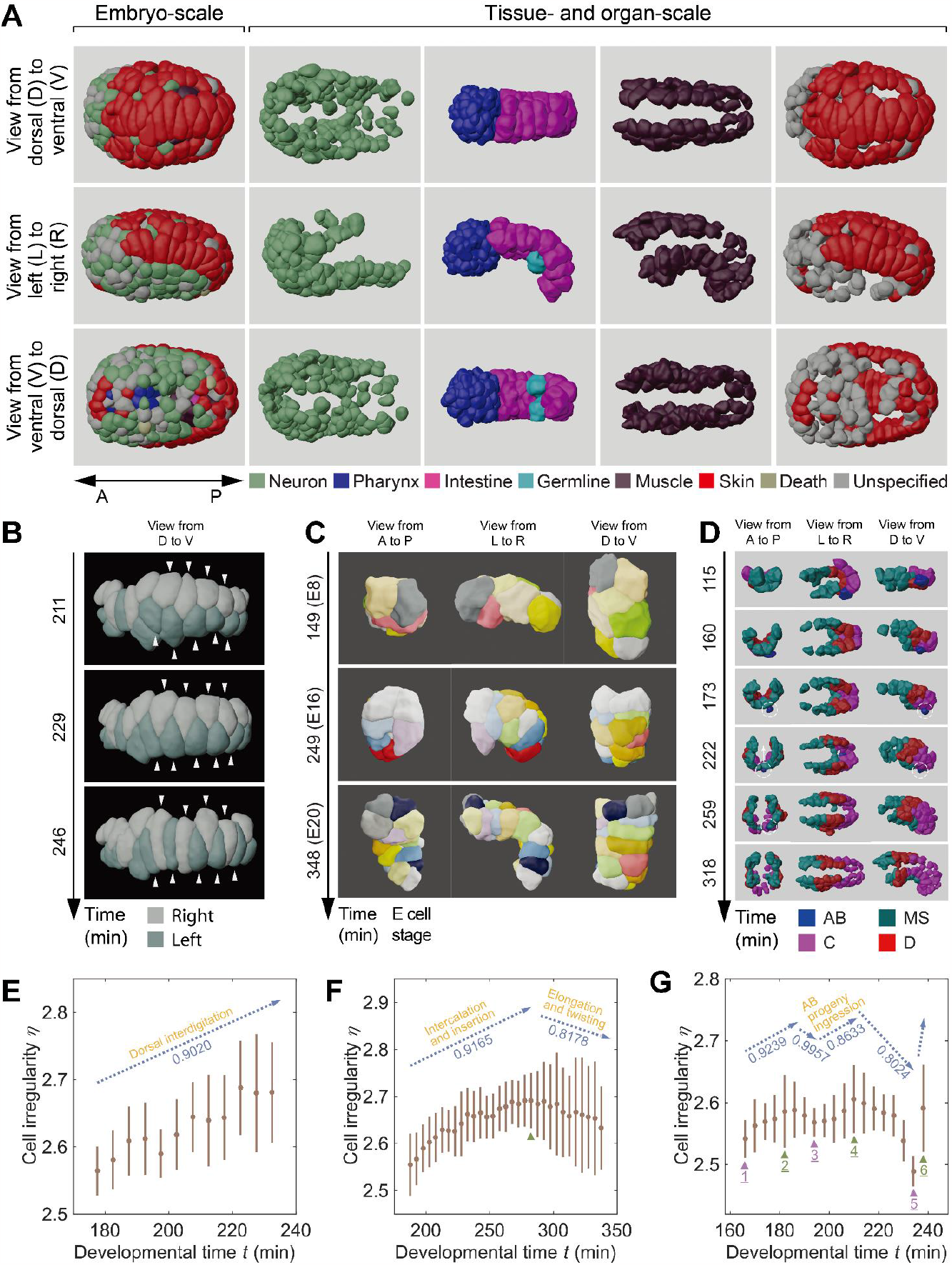
Qualitative and quantitative shape dynamics from cellular to embryonic scales. (**A**) Embryo-, tissue- and organ-scale shapes viewed from three different directions. (**B**) The cell shape changes during dorsal intercalation with the absolute time listed on the left. (**C**) The cell shape changes during intestinal morphogenesis with the absolute time listed on the left. (**D**) The cell shape changes during body wall muscle assembly with the imaging time listed on the left. Here, the ingression of the AB progeny (*i.e.*, the ABprpppppaa cell) is highlighted by a dashed white circle. (**E**) The cell irregularity (*η*; with the solid circle representing the average and the vertical line representing the standard deviation within a constant interval) dynamic of the cells participating in dorsal interdigitation over developmental time (*t*; with the last time point of the 4-cell stage as the time zero), in accordance with (**B**). Here, a total of nine neighboring cells with obvious movements toward the central line in the embryo dorsal are taken into account, as marked in (**B**); the correlation coefficient of the monotonic η – *t* curve is labeled at the top. (**F**) The cell irregularity (η; with the solid circle representing the average and the vertical line representing the standard deviation within a constant interval) dynamic of the cells participating in intestinal morphogenesis over developmental time (*t*; with the last time point of the 4-cell stage as the time zero), in accordance with (**C**). Here, all the E cells are taken into account; the global maximum of the η – *t* curve is indicated by a green triangle where the correlation coefficient of the monotonic partial curves it partitions are labeled at the top. (**G**) The cell irregularity (η; with the solid circle representing the average and the vertical line representing the standard deviation within a constant interval) dynamic of the AB progeny (*i.e.*, the ABprpppppaa cell) participating in body wall muscle assembly over developmental time (*t*; with the last time point of the 4-cell stage as the time zero), in accordance with (**D**). Here, the data of all eight *C. elegans* wild-type embryo samples are taken into account; the local maximums and minimums of the η – *t* curve are indicated by green and purple triangles respectively, where the correlation coefficient of the monotonic partial curves they partition are labeled at the top.

Our cell-resolved morphological map covers not only the morphogenetic events in the first half of *C. elegans* embryogenesis like gastrulation (Movie S6) but also the ones afterward. To further quantitatively probe into the tissue and organ morphodynamics during *C. elegans* embryogenesis, we choose three already-known events for the case study: dorsal interdigitation (Figure 3B) [Chisholm *et al., WormBook*, 2005; Altun *et al., WormAtlas*, 2009], intestinal morphogenesis (Figure 3C) [Thorpe *et al., Cell*, 1997; Asan *et al., PLoS Genet.*, 2016], and body wall muscle assembly (Figure 3D) [Altun *et al., WormAtlas*, 2009; Guan *et al., Dev. Genes Evol.*, 2020]. Here, we adopt the dimensionless surface-to-volume ratio (defined as 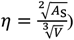) to evaluate cell irregularity, which has successfully revealed the dependence of cell irregularity on cell motility and cell lineage as reported before [Cao *et al., Nat. Commun.*, 2020]. First, for dorsal interdigitation, during which two rows of cells arranged on the two sides of the central line in the embryo dorsal move toward the central line and engage into one row, the cell irregularity rises as the cell shape is squeezed and becomes narrow (Figure 3E; Movie S7). Second, for intestinal morphogenesis, the cell irregularity shows an apparent low-high-low pattern (Figure 3F), where the former half corresponds to the intercalation and insertion of particular E cells (*i.e.*, the Ealpa, Earpa, Ealpp, Earpp cells) while the latter half corresponds to elongation and twisting (Figure S13; Movie S8). Third, for body wall muscle assembly, cells originated from different lineages (*i.e.*, AB, MS, C, D lineages) are organized concordantly in space as a whole (Figure 3D; Movie S9); a fascinating problem is how the cells originated from different lineages proceed this process in concert, in particular to the only AB cell, the ABprpppppaa cell. Revealed by our dataset, there are three reproducible pairs of peak and valley in its cell irregularity curve; it’s observed that this cell would ingress from the outside of the body wall muscle periphery, along with its cell shape changing from spherical to narrow and then returning to spherical after settling down (Figure 3G; Figure S14; Movie S10). All three cases support that the regulated morphogenetic events can be depicted by cell morphological quantification such as cell irregularity, demonstrating the potential of our morphological map to imply or predict the biological mechanisms underlying the stereotypical migration and deformation of cells.

### Cell size asymmetry between daughter cells in the anterior-posterior direction can be universally amplified by Notch signaling

With the cell-resolved morphological map of *C. elegans* embryogenesis, we aim to decipher how the cellular developmental properties are associated with each other, including but not limited to cell size (*i.e.*, cell volume and cell surface area) and cell-cell contact relationship and area (with potential for signaling transduction). A simple validated relationship is the negative correlation between cell size asymmetry and cell cycle length asymmetry (Figure S15; Table S9), in consistency with the previous theory that the content of some cell-cycle-related factors is positively correlated to the cell volume allocated from cytokinesis [Fickentscher *et al., New J. Phys.*, 2018]. Another previously studied relationship is the small cell size and apoptotic cell fate [Cordes *et al., Development*, 2006; Sethi *et al., PLoS Biol.*, 2022]; taking two *C. elegans* embryonic apoptotic cells, MSpaapp and ABprpppppa, as examples, the former proceeds apoptosis as soon as it’s produced while the latter is first involved in the development of spike tail and then proceeds apoptosis at very late stage [Sulston *et al., Dev. Biol.*, 1983]; in spite of the different onset timing for activating programmed cell death, they are both produced with an initial cell size smaller than their sister cells’ (Figure 4A), which might destine their apoptotic cell fate at the very beginning. Given these pivotal roles of cell size, next, we’ll utilize it as a center to study how cellular developmental properties are programmed together to push forward embryogenesis.

**Fig. 4.**
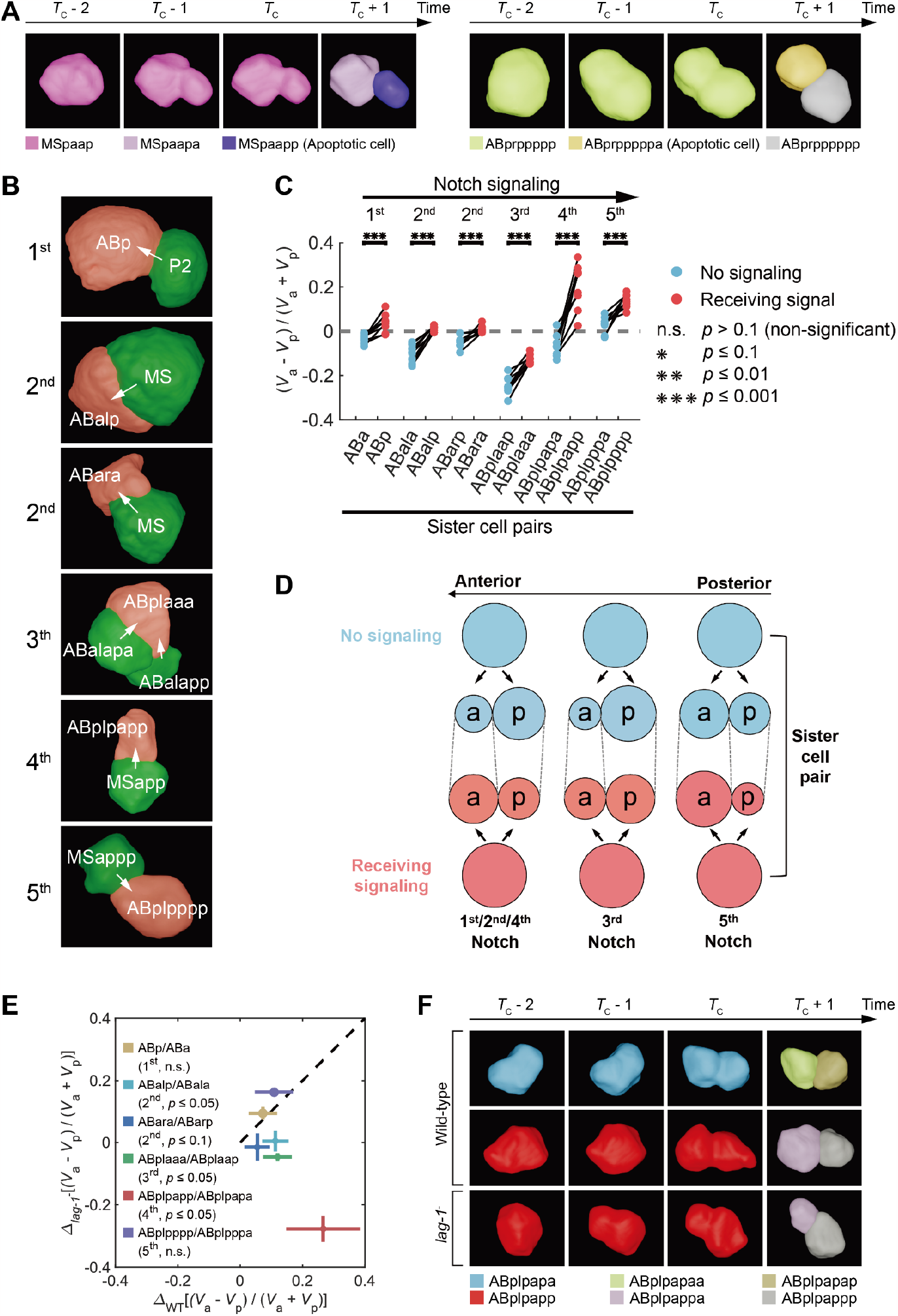
The mediation effect of Notch signaling on the cell size asymmetry between daughter cells in the anterior-posterior direction. (**A**) The morphological illustration for the asymmetric division of two representative apoptotic cells’ mothers, the MSpaap (left panel) and ABprppppp (right panel) cells. Here, *T*_*C*_ denotes the last time point when the dividing cell still has a whole cell membrane, and the morphological data at time points *T*_*C*_ − 2, *T*_*C*_ − 1, *T*_*C*_, *T*_*C*_ + 1 are illustrated together. (**B**) The morphological illustration for the cells that send and receive Notch signaling, painted in green and red respectively. (**C**) The cell volume asymmetry between daughter cells (*i.e.*,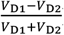) in the six sister cell pairs differentiated through Notch signaling. Here, for each of the six sister cell pairs, the cell with no signaling is painted in blue while the cell receiving signaling is painted in red; the data from all eight wild-type embryo samples are presented, where the two data points of a sister cell pair from the same embryo samples are connected with a solid black line; the statistical significance is obtained by the one-sided Wilcoxon rank-sum test (non-significant, *abbr.*, n.s., *p* > 0.1; *, *p* ≤ 0.1; **, *p* ≤ 0.01; ***, *p* ≤ 0.001) and is listed on the right. (**D**) The shift of cell volume asymmetry between sister cell pairs (*i.e.*,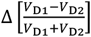) in wild-type (horizontal coordinate) and RNAi-treated (vertical coordinate) embryo samples. Here, the data from all eight wild-type embryo samples and all two *lag-1*^-^ embryo samples are presented, with the solid circle representing the average and the orthogonal lines representing the standard deviation; the statistical significance is obtained by the one-sided Wilcoxon rank-sum test and is listed on the left. (**E**) The morphological illustration for the division of the ABplpapp cell that receives the 4^th^ Notch signaling in the wild-type embryo (middle row), compared with its sister cell in the wild-type embryo (upper row) and its perturbed state in the *lag-1*^-^ embryo (lower row). Here, *T*_*C*_ denotes the last time point when the dividing cell still has a whole cell membrane, and the morphological data at time points *T*_*C*_ − 2, *T*_*C*_ − 1, *T*_*C*_, *T*_*C*_ + 1 are illustrated together. (**F**) The graphic summary for the mediation effect of Notch signaling on the cell size asymmetry between daughter cells in the anterior-posterior direction.

To elucidate how cell fate breaks its asymmetry along with other cellular developmental properties just like cell size, we focus on Notch signaling, which has been known to specify the fate of a cell from its sister cell through physical cell-cell contact [Priess, *WormBook*, 2005]; briefly, round-by-round signaling transduction would occur between groups of specific cells, where the ligand spreading on a signaling cell’s surface effectively binds the receptor on another cell’s surface but not its sister cell’s surface; such distinct contact leads a pair of sister cells to undergo differential transcriptional programs and turn into different cell fates. Here, we use our cell-resolved morphological data to illustrate the five rounds of Notch signaling events reported before, to show the contact relationship and area between cells (Figure 4B) [Chen *et al., Genetics*, 2018]. For each within a pair of sister cells, we calculate the cell size asymmetry between its two daughter cells, 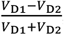, where D1 denotes its anterior daughter cell, D2 denotes its posterior daughter cell, and *V* denotes the cell volume. Strikingly, all the cells receiving signaling exhibit significantly higher cell size asymmetry between their daughter cells in the anterior-posterior direction, compared to the ones of their sister cells that don’t receive signaling (one-sided Wilcoxon rank-sum test, *p* ≤ 0.001, for all sister cell pairs) (Figure 4C). Such a significant trend is also the same with respect to cell surface area asymmetry (one-sided Wilcoxon signed-rank test, *p* ≤ 0.01 for all sister cell pairs) (Figure S16), and there’s no exception for all the sister cell pairs and all the wild-type embryo samples. This increment strongly indicates that, after a mother cell receives Notch signaling, its daughter cell produced relatively in the anterior and posterior would gain more and less cell content respectively; concerning the defaulted cell size asymmetry in the sister cells without receiving signaling, there are a total of three subtypes: 1. In the 1^st^, 2^nd^, and 4^th^ Notch signaling events, the cell size asymmetry is completely swapped; 2. In the 3^rd^ Notch, the anterior daughter cell is larger than its cousin cell without receiving signaling but is still smaller than its sister cell; 3. In the 5^th^ Notch, the anterior daughter cell becomes even larger (Figure 4D).

To further validate that such an effect is caused by Notch signaling, we generate two embryo samples with RNAi treatment that knockdowns the gene *lag-1* (the terminal effector of the Notch signaling pathway) (Table S3; Materials and Methods - RNA interference) [Priess, *WormBook*, 2005]. The difference of 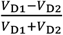 between the sister cell pair (*i.e.*, 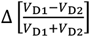, always with a positive value in a wild-type condition), in which one receives Notch signaling and the other doesn’t, is significantly changed in four out of six signaling Notch events (Figure 4E), supporting the role of the Notch signaling pathway in increasing cell size asymmetry during cell division [Chen *et al., Genetics*, 2018]; the remaining two Notch signaling events without significant effect might be due to the insufficient number of embryo samples or their weak cell size asymmetry as default, which is worth of verification in the future. Last but not least, a series of comparative morphological snapshots on the cells involved in the 4^th^ Notch signaling event in wild-type and RNAi-treated embryo samples are shown in Figure 4F, showing how the ABplpappa cell acquires larger cell size than its sister cell (*i.e.*, the ABplpappp cell), but such a situation doesn’t happen in either their cousin cells (*i.e.*, the ABplpapaa and ABplpapap cells) or themselves with the Notch signaling pathway blocked.

### Consecutive asymmetric cell divisions substantially contribute to the remarkably large size of the *C. elegans* kidney cell

Compared to the other five cells that receive Notch signaling, the ABplpapp cell in the 4^th^ Notch signaling event has the largest shift (*i.e.*, enhancement in the anterior-posterior direction) of cell size asymmetry after cell division, making its anterior daughter cell always larger than its posterior daughter cell, no matter for cell volume or cell surface area (Figure 4C; Figure S16); the change on its shift after blocking Notch signaling pathway is also the most severe, further validating the role of Notch signaling on mediating its cell size asymmetry after cell division (Figure 4E). Previous research has reported that the ABplpapp cell is specified by Notch signaling for producing the only kidney cell (also called the excretory cell), the ABplpaappaap cell (*i.e.*, great-granddaughter of the ABplpapp cell) in *C. elegans*, which is the largest cell at the adult stage with a vital function of osmotic/ionic regulation and waste elimination (Figure 5A; Figure S17) [Nelson *et al., J. Exp. Zool.*, 1984; Buechner *et al., Dev. Biol.*, 1999; Zhao *et al., J. Biol. Chem.*, 2005; Altun *et al., WormAtlas*, 2009]. Nevertheless, it’s still unclear how the kidney cell’s remarkably large size is achieved and associated with the known signaling-induced cell fate specification.

**Fig. 5.**
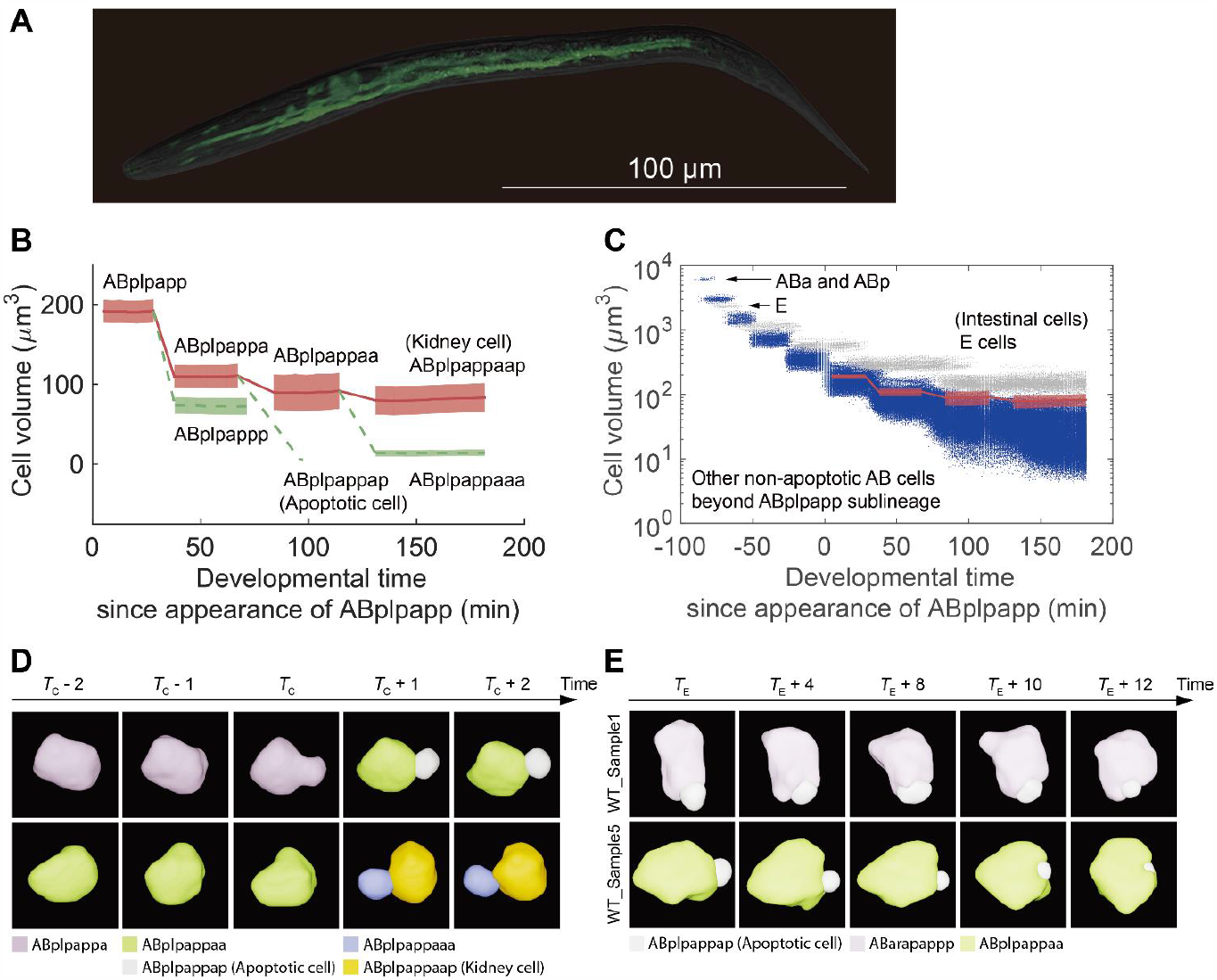
The remarkably large size of *C. elegans* kidney cell ABplpappaap contributed to by consecutive asymmetric cell divisions during embryogenesis. (**A**) The H-shaped kidney cell fluorescently labeled by GFP at the adult stage [Zhao *et al., J. Biol. Chem.*, 2005], illustrated with the merged images from confocal microscopy and differential interference contrast microscopy. (**B**) The cell volume changes over time in ABplpapp sublineage. Here, after setting the appearance of the ABplpapp cell as the time zero, for each cell, its overlapped duration among the eight wild-type embryo samples is considered, where the average and standard deviation of the eight sets of data points are illustrated by line and shade, respectively. For the illustrated cells, red and solid lines are used for labeling the ancestors of the ABplpappaap cell and itself, while green and dashed lines are used for labeling their sister cells. (**C**) The cell volume changes over time in ABplpapp sublineage as well as the other non-apoptotic AB cells and the intestinal E cells. For the additional AB and E cells for comparison, all their data points from eight wild-type embryo samples since the last time point of the 4-cell stage are plotted. (**D**) The asymmetric division of the kidney cell ABplpappaap’s ancestors, the ABplpappa (upper row) and ABplpappap (lower row) cells, illustrated with the morphological data from a *C. elegans* wild-type embryo sample. Here, *T*_*C*_ denotes the last time point when the dividing cell still has a whole cell membrane, and the morphological data at time points *T*_*C*_ − 2, *T*_*C*_ − 1, *T*_*C*_, *T*_*C*_ + 1, *T*_*C*_ + 2 are illustrated together. (**E**) The engulfment of the apoptotic cell ABplpappap by the ABarapappp (upper row) and ABplpappaa (lower row) cells, illustrated with the morphological data from the *C. elegans* wild-type embryo samples WT_Sample1 and WT_Sample5 respectively. Here, *T*_E_ denotes an arbitrarily-selected time point when the apoptotic cell is yet to be engulfed, and the morphological data at time points *T*_E_, *T*_E_ + 4, *T*_E_ + 8, *T*_E_ + 10, *T*_E_ + 12 are illustrated together.

To answer the question above, we further probe into the cell size change during kidney cell production. As the two cell size features (*i.e.*, cell volume and cell surface area) show the same asymmetry dynamics including both their relationship with cell cycle length asymmetry as well as their shift mediated by Notch signaling in the wild-type embryo, hereafter we will take the cell volume as the representative unless specified otherwise. Strikingly, in addition to the ABplpapp cell whose cell volume asymmetry is directly mediated by Notch signaling (Figure 4C; Figure 4E), its daughter ABplpappa cell and granddaughter ABplpappaa cell, namely the grandmother and mother of the kidney cell ABplpaappaap, also undergo asymmetric cell division consecutively (Figure 5B; Movie S11; Movie S12). Compared to the other non-apoptotic AB cells beyond ABplpapp sublineage, the volume of the ABplpapp cell is moderate initially and its descendants that produce the kidney cell gradually climb to the top among all non-apoptotic AB cells after three rounds of asymmetric cell divisions (Figure 5C); the declining rate of cell volume over generations is apparently smaller than that of the other non-apoptotic AB cells, leading to a continuous relative enlargement of the kidney cell and its ancestors. It is worth noting that, although it’s known that the kidney cell will become the largest cell at the adult stage, the kidney cell is still smaller than the intestinal cells (*i.e.*, E cells) by the end of *C. elegans* embryogenesis, implying that postembryonic growth goes on contributing to its remarkably large size as well (Figure 5C). Despite that the sister cells of the kidney cell and its ancestors acquire relatively small cell volume so as to increase cell volume heterogeneity, they are still capable of being programmed to have different cell fates; taking the ABplpappap and ABplpappaaa cells that have similar volumes as examples, the latter cell turns into a neuron while the former cell proceeds apoptosis, whose corpse would be randomly engulfed and digested by either of two unique cells, the ABarapappp or ABplpappaa cell (Figure 5E; Movie S11; Movie S12) [Ellis *et al., Cell*, 1986].

### Cell size asymmetry can be extensively regulated by multiple factors

As cell size asymmetry has been demonstrated to be mediated by Notch signaling and coupled with consequent cell fate specification (Figure 4C; Figure 4E), whether akin symmetric breaking is universal in more cells and regulated by more factors over embryogenesis is a profound problem. Next, we seek to dig into the potential factors that impact cell size asymmetry. First of all, we attempt to test Wnt signaling, another well-known signaling pathway responsible for *C. elegans* embryonic cell fate specification in parallel to Notch signaling [Eisenmann, *WormBook*, 2005]; for instance, under the guidance of Wnt signaling, the AB cells (the 1^st^ somatic blastomere) exhibit differential gene expression patterns along the anterior-posterior direction [Zacharias *et al., PLoS Genet.*, 2015], while the EMS cell (the 2^nd^ somatic blastomere) gives rise to mesoderm (*i.e.*, the MS cell and its offsprings) and endoderm (*i.e.*, the E cell and its offsprings) in its anterior and posterior daughter cell respectively [Thorpe *et al., Cell*, 1997]. To this end, we generate two embryo samples with RNAi treatment that knockdowns the gene *pop-1* (the terminal effector of the Wnt signaling pathway) (Table S3; Materials and Methods - RNA interference). Because the cell division orientation is severely damaged, unambiguous cell identity assignment based on the initial position of a cell right after the cytokinesis of its mother cell and with the suffix a/p/l/r/d/v (*i.e.*, anterior/posterior/left/right/dorsal/ventral) is unavailable [Sulston *et al., Dev. Biol.*, 1983]; therefore, for a pair of sister cells, we calculate their cell volume asymmetry without positional bias by calculating the absolute value of 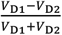(*i.e.*,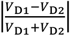). A total of 257 cell division events shared by both wild-type and RNAi-treated embryo samples in the first half of *C. elegans* embryogenesis, which have a complete lifespan recorded in the mother cell as well as the two daughter cells, are taken into account. A significant decline in cell volume asymmetry is found in the RNAi-treated embryo samples, compared to the wild-type ones (one-sided Wilcoxon rank-sum test, *p* = 1.44 × 10^−4^), with an average value changed from 0.1194 to 0.0877 (Figure 6A). Besides, the number of cells with weak volume asymmetry in their daughters (defined as 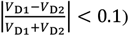) becomes larger in the RNAi-treated embryos with the Wnt signaling pathway blocked, and they have fewer cells with a strong volume asymmetry (defined as 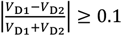) (Figure 6A). These statistical results support the role of the Wnt signaling pathway in increasing cell volume asymmetry without positional bias after cell division.

**Fig. 6.**
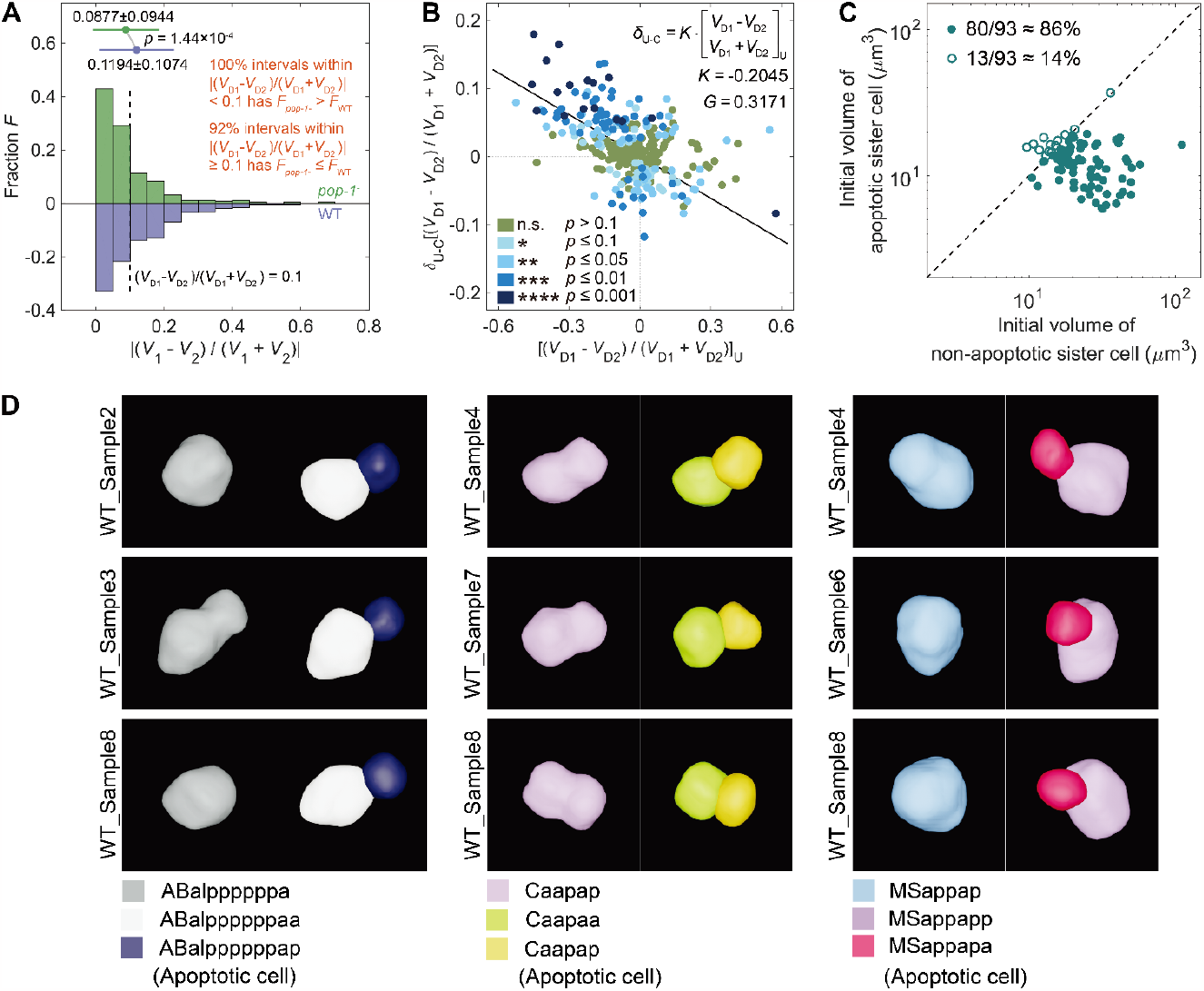
The extensive factors that can regulate cell size asymmetry between daughter cells. (**A**) The distribution of cell volume asymmetry between daughter cells without positional bias (*i.e.*,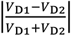) in the wild-type and *pop-1*^-^ embryos. Here, the statistical significance is obtained by the one-sided Wilcoxon rank-sum test and is listed at the top. (**B**) The negative correlation between the shift of cell volume asymmetry after mechanical compression (*i.e.*,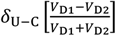) and the default in the uncompressed state (*i.e.*,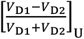). Here, the result of proportional fitting between 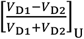 and 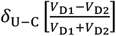 is shown with a solid line, with the proportional coefficient (*i.e., K*) and goodness of fit (*i.e., G*) listed in the top right corner; the statistical significance for the shift of cell volume asymmetry in each cell division event (presented with blue dots) is obtained by the one-sided Wilcoxon rank-sum test (non-significant, *abbr.*, n.s., *p* > 0.1; *, *p* ≤ 0.1; **, *p* ≤ 0.05; ***, *p* ≤ 0.01; ****, *p* ≤ 0.001) and is listed in the bottom left corner. (**C**) The smaller initial volume of an apoptotic sister cell (vertical coordinate) compared to the one of its non-apoptotic sister cell (horizontal coordinate). Here, the data of a total of 93 properly-recorded sister cell pairs are presented, among which 80 have a relatively smaller initial volume for their apoptotic sister cells on average. (**D**) The morphological illustration for the asymmetric divisions of three representative apoptotic cells’ mothers from the AB (left panel), C (middle panel), and MS (right panel) lineages. Here, the morphological data at the last time point when the cell still has a whole membrane, as well as the first time point when the cell division is completed, are illustrated together.

Apart from the cell volume asymmetry increased by the Wnt signaling, a previous experimental observation on *C. elegans* embryogenesis up to the 28-cell stage revealed that a certain proportion of cell division events acquire decreased cell volume asymmetry between the daughter cells when the eggshell is removed (Fickentscher *et al., Sci. Rep.*, 2017). Conversely, we wonder if external mechanical compression, which is supposed to enhance the inner pressure and change the cell positions for an embryo, would increase cell volume asymmetry. Hence, we adopt the 17 *C. elegans* wild-type embryo samples imaged under external mechanical compression from our previous paper [Cao *et al., Nat. Commun.*, 2020], where the originally ellipsoidal shape of the eggshell is deformed as an elliptical cylinder whose crosssection parallel to the shooting direction of imaging has an approximate ratio of width (9.4675 ± 0.2693 μm) to height (26.7989 ± 0.0223 μm) as 1:3 (Table S10). As those embryo samples were recorded for the first half of *C. elegans* embryogenesis (*i.e.*, roughly from the 4- to 350-cell stages), we take the 285 cells recorded with a complete lifespan for their daughter cells and themselves into account (Table S4). We calculate the shift of cell volume asymmetry (*i.e.*,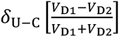) compared to the default in the mechanically uncompressed state (*i.e.*, 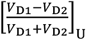); note that 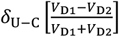 is positive if the volume of the anterior/left/dorsal daughter cell (D1) becomes relatively smaller under external mechanical compression, and *vice versa*. Intriguingly, there is a modest negative correlation between the two variables, with 151 cell division events (∼50%) exhibiting significantly changed cell volume asymmetry (one-sided Wilcoxon rank-sum test, *p* ≤ 0.1) and more than a half of them fall in the second quadrant (*i.e.*,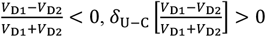) (Figure 6B;Table S11). This finding matches the previous comparison of ABpl cell’s daughter cells between mechanically uncompressed and compressed embryos [Thiels *et al., Bioinformatics*, 2021]. Our overall evaluation for multiple cells at multiple generations indicates the compensatory effect of external mechanical compression on cell volume asymmetry, against the effect of the Wnt signaling pathway.

As cell volume asymmetry is widely spread under both biochemical and biomechanical mediations (Figure 6A; Figure 6B), we’re curious if the cells with high volume asymmetry in their daughter cells are correlated with a fate specification. In our dataset, an amount of 93 sister cell pairs with one non-apoptotic cell and one apoptotic cell are recorded without data point loss at both their first appearances in at least one*C. elegans* wild-type embryo sample. It’s found that the majority of apoptotic cells (78 out of 93 sister cell pairs, >80%) acquire both smaller average volume and surface area than those of their sister cells right after the cytokinesis of their mother cells, reproducibly among embryo samples and regardless of lineal origin (Figure 6C; Figure 6D; Figure S18; Table S12); these quantitative results mean that the initial cell size asymmetry between an apoptotic cell and its sister cell, previously reported in specific population like neuroblast, is actually at work in a global manner throughout *C. elegans* embryogenesis [Cordes *et al., Development*, 2006; Sethi *et al., PLoS Biol.*, 2022].

### Customized software for visualizing and analyzing *C. elegans* embryonic morphological map

To facilitate convenient access to our cell morphological data, we take advantage of the public software *ITK-SNAP*, which can efficiently visualize a 3D image of both raw and segmented cells within three orthogonal crosssections as well as by generating rendered objects [Yushkevich *et al., EMBC*, 2016]. Based on it, we build up a new version that customizes our *C. elegans* data with multiple embryo samples, time points, cell identities, and quantitative properties (*i.e.*, cell volume, surface and contact area), not only for direct visualization but also for comprehensive analysis (Figure 7A). The new version of this software is termed *ITK-SNAP-CVE* (*C. elegans* Virtual Embryogenesis), where all the raw images and processed results of eight mechanically uncompressed wild-type embryo samples and four mechanically uncompressed RNAi-treated embryo samples first presented in this work, along with seventeen mechanically compressed wild-type embryo samples published in our previous paper, are provided in designated and consistent formats (Table S13; Materials and Methods - Systematic check for data consistency). There are different display modes to visualize the cells, including “*Show all cells*” (Figure 7B), “*Show master cells only*” (Figure 7C), “*Show master cells and neighbours*” (Figure 7D), and “*Show master cells and other cells*” (Figure 7E); here, the master cells are the ones arbitrarily chosen by the user through inputting or ticking cell name (*i.e.*, cell identity) with any preference of cell lineage (Figure 7F) and cell fate (Figure 7G), while the other cells beyond them can be shown with tunable opacity. In addition to the main menu (placed in the top right of the interface) for selecting display modes, embryo samples, time points, master cells, and other cells’ opacity, a submenu (placed in the bottom right of the interface) is also designed for tracking any single cell of interest with its volume, surface and contact area. To sum up, such an integrative tool is anticipated to allow researchers to navigate the cell-resolved *C. elegans* embryonic morphological map freely and facilitate the visualization and analysis of this informative data, effectively supporting relevant research in all kinds of topics.

**Fig. 7.**
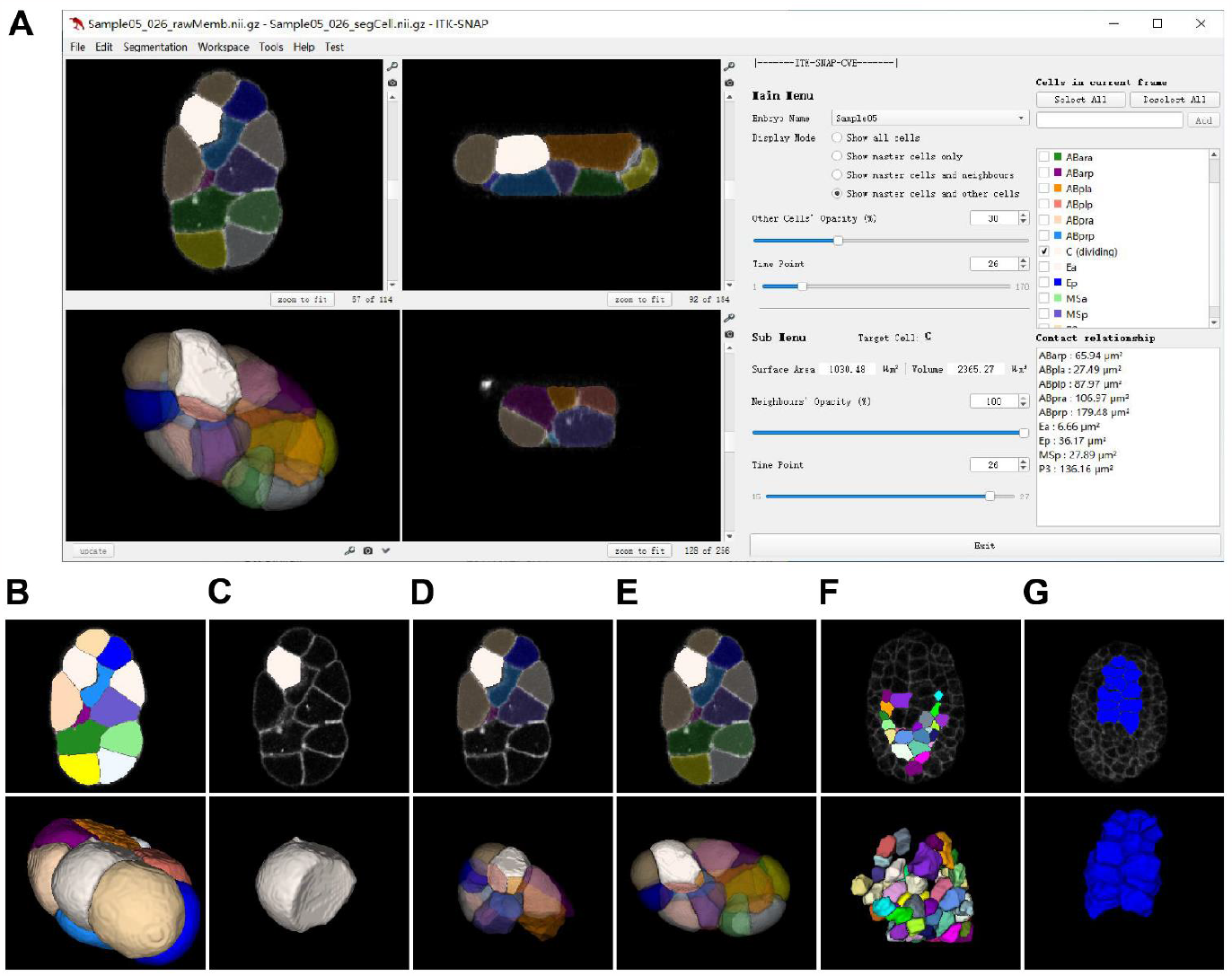
The software *ITK-SNAP-CVE* customized for visualizing and analyzing the *C. elegans* embryonic morphological map. (**A**) The graphical user interface. (**B**) The 2D and 3D snapshots for all the cells in an embryo, outputted by the display mode “*Show all cells*”. (**C**) The 2D and 3D snapshots for a specific cell in an embryo, outputted by the display mode “*Show master cells only*” and exemplified by the C cell. (**D**) The 2D and 3D snapshots for a specific cell and all its neighbours in an embryo, outputted by the display mode “*Show master cells and neighbours*” and exemplified by the C cell. (**E**) The 2D and 3D snapshots for a specific cell and all other cells in an embryo, outputted by the display mode “*Show master cells and other cells*” and exemplified by the C cell. (**F**) The 2D and 3D snapshots for all the cells from the same lineage, exemplified by the MS lineage. (**G**) The 2D and 3D snapshots for all the cells with the same fate, exemplified by the intestinal cells (*i.e.*, E cells).

### Concluding remarks

In this work, we set up a comprehensive system for reconstructing and studying the *C. elegans* embryonic morphological map at the single-cell resolution, which is comprised of a novel transgenic strain, a new algorithm *CMap* for segmenting and processing 3D cell images, and customized software *ITK-SNAP-CVE* with strict data quality control (Figure 1A; Figure 1B; Figure 7A). The morphological map covers almost all the cells that appear in *C. elegans* embryogenesis (Table S4), with their cell shape, volume, surface and contact area measured quantitatively, as well as their cell identity, lineage, and fate (Figure 2; Figure 3). The direct output information on cell volume and cell surface area is used to explore the cell size programming associated with Notch signaling, Wnt signaling, mechanical compression, and cell fate (Figure 4, Figure 5, Figure 6); the indirect shape information on cell irregularity (calculated with cell volume and cell surface area) is used to distinctly characterize the typical morphogenetic events including dorsal interdigitation (Figure 3E), intestinal morphogenesis (Figure 3F), and body wall muscle assembly (Figure 3G). These discoveries together demonstrate the value of this abundant cell morphological data for biological research concerning different dimensions (*e.g.*, cell shape, volume, surface and contact area), events (*e.g.*, morphogenetic developmental landmarks), and conditions (*e.g.*, RNAi treatment and mechanical compression).

Although the proposed system exhibits considerable performance in reconstructing cell morphology during *C. elegans* embryogenesis, there is still a total data point loss rate of nearly 8%. Estimated with the (data point) loss rate of a given 3D image, for temporal dimension, the loss rate increases gradually over embryogenesis (Table S7); for spatial dimension, the loss is the most severe on the edge of the embryo: among the cells with different fates, the skin cells exhibit the highest loss rate, while among the cells from different lineages, the C cells (*i.e.*, major contributors to skin), exhibit the highest loss rate (Figure S19; Table S8). The failure in recognizing and segmenting membrane fluorescence is caused by both experimental and computational reasons as follows. 1. Unlike the cell-cell contact boundary composed of two layers of cell membranes with fluorescence from neighboring cells, the cell boundary on the edge of the embryo and exposed to the external environment has only one layer of cell membrane with fluorescence, which is supposed to be 50% weaker (Figure S20); since the extra-embryonic space is regarded as a dilatable region in *CMap* algorithm just like the one of a cell, it would invade the embryo when the membrane fluorescence on the edge of the embryo is not enough. 2. The attenuation of laser intensity through the embryo body makes the membrane fluorescence in the top focal planes weaker than the one in the bottom focal planes (Figure S20). 3. Cell apoptosis undergoes cell size shrinkage, where the membrane fluorescence acquires an enhanced thickness and gets blurred (Figure S20). Figure S21 shows how different laser intensities during imaging impact the fluorescence distinctness. All the shortcomings above echo the future improvement in various aspects (*e.g.*, strain, microscopy, algorithm) to realize better performance in cell morphology reconstruction [Guan *et al., Comput. Struct. Biotechnol. J.*, 2022].

The cell morphological data generated by our system is multidimensional, consisting of information on cell shape, cell volume, cell surface and contact area, cell position, cell identity, cell lineage, cell fate, and so on. It serves as a valuable resource to investigate how those cellular developmental properties interact with each other and how they respond to perturbation like RNAi on specific genes and mechanical compression (Figure 4; Figure 5; Figure 6). Systematic analysis considering the factors listed above has revealed multimodal mechanisms for cell size programming, however, identifying the association and/or dependence is only the first step, which evokes both upstream and downstream research around this problem. About the upstream, it’s still unknown how the Notch and Wnt signaling influences the cell size asymmetry; in other words, the complete pool of functional genes linking the signaling pathways to cytoskeleton dynamics remains to be identified [Ajduk *et al., Mol. Hum. Reprod.*, 2016]. Also, how does the mechanical compression influence the cell size asymmetry across multiple cells exactly; perhaps it reflects the difference in intercellular pressure [Thiels *et al., Bioinformatics*, 2021; Cuvelier *et al., Biophys. J.*, 2023]? About the downstream, as the cell size asymmetry is accurately programmed with a limited tolerance for successful embryogenesis and is critical for cell cycle length asymmetry as well as consequential cell arrangement progression [Jankele *et al., eLife*, 2021; Guan *et al., Phys. Rev. E*, 2021; Kuang *et al., PLoS Comput. Biol.*, 2022], it’s elusive why significantly shifted cell size asymmetry caused by mechanical compression doesn’t induce lethality; will it expose any compensatory or fail-safe mechanism [Jelier *et al., Cell Syst.*, 2016]? These and other scientific questions can be explored in depth using the *C. elegans* embryonic morphological map established in this work.

## METHOD SUMMARY

All methods used in this study are recapitulated in detail in the supplementary materials, which include the following six sections: (i) methods summary, (ii) supplemental figures, (iii) captions for supplemental tables, (iv) captions for supplemental movies, (v) references and notes, (vi) author contributions.

## ACKNOWLEDGMENTS

We thank all the members of Yan Lab, Tang Lab, and Zhao Lab for the fruitful discussion and constructive comments. We appreciate Dr. David Smith and Yixuan Chen for their assistance in improving the paper materials and Ahjol Hyrat for preliminary analysis on *C. elegans* developmental symmetry breaking. We are grateful to Prof. Zhuo Du, Dr. Xuehua Ma, and Dr. Zhiguang Zhao for providing the information on *C. elegans* cell identity and cell fate. Gratitude is extended to KC Yung for his help in the construction of *ITK-SNAP-CVE* and Datatang Technology Inc. for manual annotation on cell morphology.

## FUNDING

This work was supported by the National Natural Science Foundation of China (12090053, 32088101) to G.G. and C.T.; by the General Research Funds (N_HKBU201/18, HKBU12101520, HKBU12101522) from the Hong Kong Research Grants Council and State Key Laboratory of Environmental and Biological Analysis grant, Hong Kong Innovation and Technology Fund (GHP/176/21SZ), and Initiation Grant for Faculty Niche Research Areas (RC-FNRA-IG/21–22/SCI/02) from Hong Kong Baptist University to Y.M., M.-K.W., L.-Y.C., and Z.Z.; and by the Hong Kong Innovation and Technology Commission (InnoHK Project CIMDA) and Hong Kong Research Grants Council (11204821) to Z.L., J.C., and H.Y.

## Author contributions

Author contributions are listed in detail in the supplementary materials.

## Data and materials availability

The data and software are available upon reasonable request and will be uploaded to the public depository before journal publication.

## SUPPLEMENTARY MATERIALS

### (i) Methods summary

#### Three-dimensional illustration for segmented cell morphologies

All the 3D illustrations, including both figures and movies, are rendered with the software *Fiji* and *Blender* [Schindelin *et al., Nat. Methods*, 2012; Blender Online Community, *Blender Foundation*, 2023].

#### Worm strains and maintenance

All the animals were maintained on NGM (Nematode Growth Media) plates seeded with *Escherichia coli* OP50 at a constant ambient temperature of 20°C . A cell nucleus and cell membrane dual-labeled strain, ZZY0535 [Chen *et al., Genetics*, 2018], was crossed with another bombardment-generated dual-labeled strain, ZZY0855, zzyIs139 [Phis-72::PH(PLC1delta1)::mCherry::*pie-1* 3’ UTR+*unc-119*(+)], to enhance the fluorescence signal intensity of cell membrane. The four markers in these two strains were rendered homozygous in a novel strain, ZZY0861. The resulting strain can ubiquitously express a fusion between histone (in the cell nucleus) and GFP (Green Fluorescent Protein), allowing nucleus identification as well as automated tracing and lineaging; meanwhile, the mCherry labeling cell membrane enables membrane segmentation. Additionally, a strain that stably expresses GFP in the kidney cell throughout L1, L2, L3, L4, and adult stages, BC10210, *dpy-5*(e907), sIs10089 [*dpy-5*(+)+rCes-*pgp-12*-GFP+pCes361], was used for exhibiting its outstandingly large cell size [Zhao *et al., J. Biol. Chem.*, 2005].

#### Fluorescence microscopy for embryonic stages

The approaches are as described before [Cao *et al., Nat. Commun.*, 2020], except for the new experimental operations and imaging parameters (*incl.*, layer number and laser intensity) designed to maintain the natural shape of the embryo’s eggshell without lateral mechanical compression: (1) For the RNAi-treated worms, 1- to 4-celled embryos were retrieved from the adults that had been subjected to injection for at least 12 hours but no longer than 24 hours, while the wild-type ones were retrieved directly without such special procedure; (2) The embryos were mounted on a polylysine-pretreated glass slide for imaging; (3) Vaseline was dotted outside the four corners of the polylysine pad, and 12 μL of Boyd’s buffer was added on top of the embryos and pad; (4) The coverslip was gently placed on top of the buffer, and the four corners of the coverslip were slowly pressed until they touched the buffer; (5) After ensuring that the embryos were within the area where the coverslip touched the buffer, all four edges of the coverslip were sealed with mol30aselineline.

Imaging was performed with the Stellaris confocal microscopy system (Leica) at a constant ambient temperature of 20°C. Images were acquired from both GFP and mCherry channels with a frame size of 712×512 pixels (resolution: 0.09 μm) and a scanning speed of 8000 Hz, using a water immersion objective. The excitation laser beams used for GFP and mCherry are 488 and 594 nm, respectively. Fluorescence images from 92 focal planes were consecutively collected for three embryos per imaging session, with a resolution of 0.42 μm from top to bottom of the embryo for each time point, which was at approximately less than 1.5-minute intervals (Table S3). Images were continuously collected for at least a total of 240 time points, during which the cell count would reach up to the 550-cell stage in a wild-type embryo and up to the 330-cell stage in an RNAi-treated embryo. The entire imaging duration was divided into five time blocks based on the given time points, *i.e.*, 1-60, 61-130, 131-160, 161-200, and 201-240. The *z*-axis compensation was 0.5-3% for the 488 nm laser and 20-95% for the 594 nm laser. The pinhole sizes for the four blocks were 2.3, 2.0, 1.6, 1.6, and 1.3 AU (Airy Unit), respectively. Prior to image analysis, all images were subjected to deconvolution, then each image was renamed sequentially.

#### Fluorescence microscopy for postembryonic stages

Images were acquired using the SP5 II or Stellaris confocal microscopy system (Leica) and analyzed using the software *LAS X* (Leica) and *ImageJ* [Abramoff *et al., Biophotonics Int.*, 2004]. Micrographs of larvae and gonads were acquired with tile scanning using the same confocal microscope as that for the embryo. Dissected gonads or intact adults were mounted with Boyd’s buffer/methyl cellulose [Murray *et al., Nat. Protoc.*, 2006] for imaging. Animals were mounted on a 1% agarose pad with 0.1 M (Molar) sodium azide in M9 buffer for imaging with a scanning speed of 200-400 Hz depending on the size of the animals. For the acquisition of 3D image stacks, imaging settings were the same as those used for the embryo, except that the *z*-resolution is 1 µm for the adult and 0.71 µm for the embryo.

#### Automated cell nucleus tracing and lineaging

The cell nuclei images (from the GFP channel) were automatically identified, traced, and lineaged using the software *StarryNite* and then visualized using the software *AceTree* [Bao *et al., Proc. Natl. Acad. Sci. U. S. A.*, 2006; Murray *et al., Nat. Protoc.*, 2006; Katzman *et al., BMC Bioinform.*, 2018]. The automated tracing and lineaging results were manually corrected up to beyond the 550-cell stage, as the full lifespans of the C lineage had to be recorded. The nucleus information, including spatial position and cell identity at each time point, was outputted in a separate file.

#### Calculation of cell volume, cell surface area, and cell-cell contact area

For the segmented 3D cell regions in *C. elegans* embryos, the cell volume (*i.e., V*) was calculated by summing the volumes of the corresponding pixels. Moreover, the alpha shape model, which is commonly used to create a 3D triangular mesh from the surface of a 3D object, was employed to extract the surface area (*i.e., A*S) of a cell and the contact area (*A*C) between two cells [Still *et al., Microsc. Microanal.*, 2020]. The detailed procedure is as follows: (1) Dilation with a thickness of a pixel added was executed on each 3D cell region, followed by the generation of a 3D triangular mesh from the dilated surface; (2) The cell surface area was calculated by summing the areas of all the triangles on the mesh; (3) The pixels contacted and between any two cell regions were detected and recorded; (4) For each of a pair of contacted cells, the partial area of its surface mesh that contains those pixels was calculated; (5) About two cells that contact each other, the larger value of the abovementioned partial area was adopted as the contact area between them. The quantification of the three morphological properties was jointly validated by two evaluations on each cell region of the eight wild-type embryos and four RNAi-treated embryos presented in this work: (1) The dimensionless cell irregularity (defined as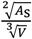) is always larger than the theoretically minimum 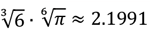in a perfect sphere; (2) The sum of the contact areas with neighboring cells never exceeds the surface area with a relative discrepancy of 20% [Cao *et al., Nat. Commun.*, 2020].

#### Automatic segmentation

In this paper, we proposed an integrated method, *CMap*, for automatically segmenting 3D fluorescence images, also known as cellular volumetric segmentation. The proposed method includes three successive parts: (1) Interpolation is performed for the 2D TIFF images collected experimentally, transforming them into 3D NIfTI images with designed sizes and equal resolutions in all the *x, y*, and *z* directions; (2) A deep 3D neural network, EDT-DMFNet (Euclidean Distance Transform Dilated Multi-Fiber Network), predicts the probability that a single pixel belongs to cell membrane; (3) With the probability map, the segmented instances (*i.e.*, multiple separate 3D cell regions) are generated. The details of these three parts are introduced below.

##### 1. Image preprocessing

For every embryo, the fluorescence images at each time point were originally captured along the *z* axis (*i.e.*, the depth direction perpendicular to the focal plane) as a stack of 2D images, namely, a 3D image. Its corresponding pixel number along *x* (*i.e.*, the width of the focal plane), *y* (*i.e.*, the height of the focal plane), and *z* axes is 715, 512, and 92, respectively. For each raw 3D image, the pixels whose fluorescence intensity is within the highest and lowest 1% of the global range were discarded. Then, the 3D image underwent spatial resampling with downsampling along the *x* and *y* axes from 712×512 to 356×256 and upsampling along the *z* axis from 92 to 214, using bilinear interpolation.

##### 2. Euclidean distance map prediction

The volumes of the rescaled 3D image were processed by EDT-DMFNet (Euclidean Distance Transform Dilated Multi-Fiber Network), which follows the structure of U-Net [Ronneberger *et al., MICCAI*, 2015], and then were transformed into a membrane-centered 3D Euclidean distance map. To increase the perceptive field within which the communications between convoluted channels are made in our small-scale network (a tiny fully connected network), inspired by DMFNet [Chen *et al., MICCAI*, 2019], we deployed a 3D dilated regression network EDT-DMFNet to incorporate the information across multiple neighboring images in different convoluted channels with high computational speed (Figure S2; Table S1). EDT-DMFNet utilized weighted fully-dilated convolution to summarize the features at different scales adaptively. While group convolution was used to implement channel-wise convolution for a small network, a multiple-stage multiplexer, which is composed of several 1×1×1 filters, was applied to route the information among groups.

EDT-DMFNet transforms the segmentation task from a pixel classification problem to a pixel probability map regression problem. The designed network output, namely the inference on the probability map of pixels belonging to cell membrane *P* ∈ [0,1]^*W*×*H*×*D*^ according to the input image *I*_0_ ∈ [0,255]^*W*×*H*×*D*^, allows multiple cell objects segmentation with small computational resource and high speed; here, *W, H*, and *D* denote the width (along *x* axis), height (along *y* axis), and depth (along *z* axis) of the 3D image, while the corresponding values in this work are 356, 256, and 214, respectively. Therefore, the *Softmax* activation function at the output layer was replaced with the *Sigmoid* activation function for probability prediction. An adaptive membrane-centered weighted loss *L* was used to measure the difference between the target map (*i.e.*, the ground-truth data) *I* and the predicted probability map Î. The loss *L* of a 3D volumetric image was calculated by taking the weighted average of the mean squared error between the target map *I* and the probability mapÎ. Here, the adaptive membrane-centered weighted mask *W*_mask_ is defined as *W*_mask_ = *μ* · *I* + *avg*{*I*}, where *μ* is a constant (0.2) that scales the probability map Î, and *avg*{*I*} is the average value of *I* over the entire 3D image. In training, the pixels of the predicted probability map Î nearby cell membrane become more significant with the weighted mean squared error (*abbr.*, MSE). Thus, the weighted mask substantially enhanced the contributions of non-zero pixels in the loss function because the non-zero value represents the probability of a pixel belonging to cell membrane in the predicted probability map *Î*. Meanwhile, the pixels of the target map *I* surrounding the cell membrane should accordingly generate larger values in the weighted mask *W*_mask_. As a result, the loss grows rapidly if the EDT-DMFNet makes the wrong prediction for the pixels near cell membrane, effectively encouraging the network to focus on the cell membrane and its surrounding pixels. In training, the input image *I*_0_ was imposed with random noises, cropped into a 128×128×128 volume, and flipped, which realized the augmentation of the training data and ensured the robustness of the network (Materials and Methods - Training data augmentation). The Adam optimizer was employed to update the network with an initial learning rate of 5 × 10^−3^ and a weight decay rate of 1 × 10^−5^, using AMSGrad gradient descent optimization. The model was trained for 50 epochs with a batch size of 8 on an NVIDIA 2080 Ti GPU. The output of EDT-DMFNet, Î, is a probability map as well as an Euclidean distance map for a 3D volumetric image, which will be subjected to the subsequent multiple cell objects segmentation.

##### 3. Cell region generation

Based on the nucleus marker-seeded watershed algorithm [Walt *et al., PeerJ*, 2014; Khan *et al., Microsc. Res. Tech.*, 2019], *CMap* pre-inserts the experimentally-measured cell nuclear positions into the Euclidean distance map Î. This strategy confers solid cell position information to cell membrane segmentation as guidance cues. The novel cell nucleus marker-based watershed algorithm successfully improved the segmentation performance of *CMap* for living *C. elegans* embryos, avoiding under- and over-segmentation problems, especially when the embryo is at the second half of its embryogenesis with more than 350 cells. Such panoramic cell segmentation was executed for all the time points recorded in every embryo.

#### Training data augmentation

Due to the rapidly changing cell morphology during embryogenesis and the uneven distribution of fluorescence signal, the 3D images captured in the experiment always have unexpected noise and indistinct intensity, which are harmful to cell membrane recognition (semantic segmentation) correctness to varying degrees. Importantly, by adding the low convoluted feature images into the upsampling layers, U-shaped network accepts spatially operated images as different training data, which allows the deep convolutional neural networks to run effectively on a small medical or biological training dataset [Ronneberger *et al., MICCAI*, 2015]. Thus, data augmentation is critical for the training of EDT-DMFNet. Appropriate and valid augmentation not only guarantees the robustness of the network but also improves its universality, avoiding the potential over-fitting and under-fitting. Because of the small amount of training data (a total of 54 3D images reconstructed with two embryo samples), the augmentation part helps to improve the extent to which the training dataset simulates real-world data [Cao *et al., Nat. Commun.*, 2020]. By perturbing the pixels’ intensity and randomly flipping and cropping an image, the 54 3D images in the training dataset were augmented to 21600 effective training 128×128×128 cube images, which significantly improves the robustness of the trained EDT-DMFNet for other wild-type embryonic images. The protocols for data augmentation in training are described below.

Random intensity scale change: the intensity of all pixels in each training image was scaled with a uniform distribution, the half-open interval [1,1.1).

Random intensity shift change: the intensity of all pixels in each training image was shifted with a uniform distribution, the half-open interval [0,0.1).

Random flip: each training image had a 50% chance to be flipped along the *x, y*, and *z* axes, respectively.

Random crop: each training cube image was cropped randomly as [128,128,128] from the original size [205,285,134].

#### Manual annotation

The ground truth of cell morphology is necessary for evaluating the performance of machine segmentation algorithms. In this work, the cell morphology of two embryo samples (“WT_Sample1” and “WT_Sample7”) was annotated by ten well-trained experts based on the segmentation results from *CMap* and the raw fluorescence images. The segmentation results were strictly checked and corrected slice by slice and cell by cell, ensuring the correctness of the ground truth established. Specifically, we gained two sets of ground truth data in different dimensions. First, the middle slice at each imaging time point throughout embryogenesis (an amount of 255 2D images for “WT_Sample1” and 205 2D images for “WT_Sample7”) was annotated for 2D comparison, providing the crosssection of 30509 cell regions in total. Second, the whole 3D stack within 100±5 -, 200±5 -, 300±5 -, 400±5 -, 500±5 -, and 550±5 -cell stages (an amount of 6 3D images for either “WT_Sample1” or “WT_Sample7”) were annotated for 3D comparison, providing the full morphology of 4046 cell regions in total.

### Performance evaluation of *CMap* and other existing cell segmentation algorithms

To evaluate the performance of *CMap* and other existing cell segmentation algorithms, including *CShaper* [Cao *et al., Nat. Commun.*, 2020], *3DCellSeg* [Wang *et al., Sci. Rep.*, 2022], *CellPose3D* [Eschweiler *et al., ICIP*, 2022], *StarDist3D* [Weigert *et al., WACV*, 2020], and *VNet* [Milletari *et al., 3DV*, 2016], we collected a total of 54 training and 12 evaluating 3D images (Materials and Meth–ds - Manual annotation) [Cao *et al., Nat. Commun.*, 2020]; note that *VNet* is a semantic segmentation network so we integrated the instance segmentation part of *CShaper* into its binary output for subsequent performance evaluation. All six algorithms were trained through the same pipeline (Materials and Methods - *Automatic segmentation*).

Each algorithm generated multiple cell objects for 12 3D images (embryo sample WT_Sample1 at 90/123/132/166/178/185 time points with 101/201/300/400/495/551 cells respectively and embryo sample WT_Sample7 at 78/114/123/157/172/181 time points with 100/203/304/400/505/552 cells respectively) (Materials and Methods - Manual annotation). Then the segmentation outputs were quantitatively evaluated with the manually annotated ground truth by two metrics, Hausdorff distance and Dice score (Figure S5). To avoid the impact from extreme cases and achieve a fair comparison, for the Hausdorff distance, we excluded the data values smaller than 1 and larger than 20; for the Dice score, we excluded the data values smaller than 0.1 and larger than 0.9. The detailed results of each 3D image and each algorithm are listed in Table S2.

### Quality control on embryo imaging and cell tracking

Every *C. elegans* wild-type embryo sample was consecutively imaged for a duration satisfying the requirements below:

1. The last moment of the 4-cell stage has to be recorded.
2. The whole C lineage has to be generated, namely, with 8 progenies in Caa sublineage, 8 progenies in Cpa sublineage, 16 progenies in Cap sublineage, and 16 progenies in Cpp sublineage. Lastly, there are 48 C cells in total, each of which has at least 5 time points recorded.
3. The final recognized cell number must be over 570 and the total edited time points must be no less than 185 (≈ 265 minutes).

With the 3D time-lapse images, all the cells without programmed death have to be tracked until their division or the last edited time point, while the apoptotic cells are tracked to their last time point when they are still distinguishable to the naked eye. The results of cell tracking were subjected to manual editing according to the procedures:

1. For each cell with a complete lifespan recorded in all embryo samples, its normalized cell cycle lengths in all embryos are noted as [*l*1, *l*2, …, *l*8]. The top 10% cells with the largest max{[*l*1, *l*2, …, *l*8]}/mean{[*l*1, *l*2, …, *l*8]} and the bottom 10% cells with the smallest min{[*l*1, *l*2, …, *l*8]}/mean{[*l*1, *l*2, …, *l*8]} would be subjected to manual check.
2. For each 3D region marked by cell identity *c*, embryo sample *j*, and time point *T*, we characterized its volume dynamics with three characteristics:
  - The relative difference in volume between daughter cells and their mother is defined as

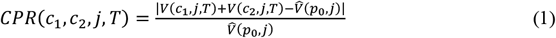

where *V*(*c*_1_, *j, t*) and *V*(*c*_2_, *j, t*) are the volumes of two daughter cells *c*_1_ and *c*_2_ at time point 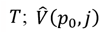 is the median volume of the mother cell *p*_0_ throughout its lifespan.
  - The volume inconsistency of cell *c* at two consecutive time points *T* + 1 and *T* is defined as

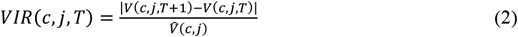
  - The volume variation of cell *c* at a specific time point *T* is defined as

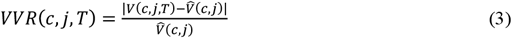

For every embryo sample, if a specific cell at a specific time point belongs to the top 1/30 ≈ 3.33% outliers for all three criteria, it would be subjected to manual check.

#### RNA interference

Gene knockdown was performed by RNAi (RNA interference) through microinjection. Primers for the amplification of the dsRNA (double-stranded RNA) template were selected based on the previous criteria [Sönnichsen *et al., Nature*, 2005; Green *et al., Cell*, 2011]. For dsRNA production, the T7 promoter was included at the 5′ ends of both forward and reverse. The primers are as follows: *lag-1*: forward: TAATACGACTCACTATAGGG TCAGTCTCTTGCAAACCACG; reverse: TAATACGACTCACTATAGGG ATGCTGCAATCGAGAATGA; *pop-1*: forward: TAATACGACTCACTATAGG TTCCCAGGAAAGTTAGGCAA; reverse: TAATACGACTCACTATAGG AAACCGACACCCGTATGAAG. PCR (Polymerase Chain Reaction) was performed using *C. elegans* N2 genomic DNA as a template in 20 μL volume using ExTaq DNA polymerase (Cat# RR001Q, TaKaRa). After checking the PCR product on 1% agarose gel, 1 μL of the PCR product was used as a template for dsRNA production with NEB HiScribe T7 Quick High Yield RNA Synthesis kit according to the manufactrer’s description. The reaction mixture was incubated at 75°C for 15 minutes in a water bath followed by turning off the heating power and incubating overnight in the same water bath for annealing of dsRNAs. The dsRNA was diluted to a concentration of 300 ng/μL in TE buffer for microinjection. At last, the automated tracing and lineaging results were manually corrected up to the 500-cell stage for two embryo samples with knockdown on *lag-1* (the terminal effector of the Notch signaling pathway) and up to the 330-cell stage for two embryo samples with knockdown on *pop-1* (the terminal effector of the Wnt signaling pathway) (Table S3).

#### Data filtering based on abnormal cell volume and cell shape

As some 3D cell regions segmented by *CMap* might be of poor quality, we devised a pipeline to filter them out from the last time point of the 4-cell stage to the last edited time point for every embryo sample. A total of six strategies were applied as listed below.

- The volume (*i.e., V*) of the cell regions was used as a criterion, which satisfies a bimodal distribution, where the extremely small cell volume is shown to be resulting from the cell region that fails to dilate toward the distinguishable cell membrane and is close to the area of the cell nucleus (Figure S6A; Figure S8). A cell region would be filtered out if it has a volume on the left of the valley of the bimodal distribution.
- The relative difference in volume between daughter cells and their mother (*i.e., CPR*) was used as a criterion. A cell region would be filtered out if it has a value over 99% of cell regions in the first half of *C. elegans* embryogenesis recorded by *CShaper* dataset (Figure S6B; Figure S7) [Cao *et al., Nat. Commun.*, 2020].
- The volume inconsistency of cell *c* at two consecutive time points *T* + 1 and *T* (*i.e., VIR*) was used as a criterion. A cell region would be filtered out if it has a value over 99% of cell regions in the first half of *C. elegans* embryogenesis recorded by *CShaper* dataset (Figure S6C; Figure S7) [Cao *et al., Nat. Commun.*, 2020].
- The volume variation of cell *c* at a specific time point *T* (*i.e., VVR*) was used as a criterion. A cell region would be filtered out if it has a value over 99% of cell regions in the first half of *C. elegans* embryogenesis recorded by *CShaper* dataset (Figure S6D; Figure S7) [Cao *et al., Nat. Commun.*, 2020].
- The existence of separate regions (*i.e., SR*), which disobeys the space continuity of cell shape, was used as a criterion. A cell region would be filtered out if it contains two cell nuclei inside but either of the middle point and the two quarter points between them doesn’t belong to the cell region (Figure S6E; Figure S8).
- The filter rate of lifespan recorded (*i.e., FRL*) was used as a criterion. A cell region would be filtered out if over half of the time points within the lifespan of this cell have been filtered out by the five criteria above (Figure S6E; Figure S8).

#### Systematic check for data consistency

To ensure that the massive data brought to the public is in the correct format, we wrote independent code to check its different aspects, including but not limited to:

- Every cell recorded in “SegCell\Embryo_Time_segCell.nii.gz” is included in “Embryo_lifescycle.csv” and “TPCell\Embryo_Time_cells.txt”.
- Every cell recorded in “SegCell\Embryo_Time_segCell.nii.gz” has been recorded in “name_dictionary.csv” and has a volume recorded in “Embryo_volume.csv” and a surface area recorded in “Embryo_surface.csv”.
- Every cell recorded in “SegCell\Embryo_Time_segCell.nii.gz” has a volume and a surface area recorded in “Embryo_Time_nucLoc.csv”.
- Every cell with a volume recorded in “Embryo_volume.csv” and a surface area recorded in “Embryo_surface.csv” has a corresponding segmented region recorded in “SegCell\Embryo_Time_segCell.nii.gz”.
- Every cell with a volume and a surface area recorded in “Embryo_Time_nucLoc.csv” has a corresponding segmented region recorded in “SegCell\Embryo_Time_segCell.nii.gz”. The cell in “Embryo_Time_nucLoc.csv” without a volume and a surface area recorded has a nucleus recorded in “Embryo_CD.csv”.
- Every cell nucleus at a specific time point recorded in “Embryo_CD.csv” are also recorded in “Embryo_Time_nucLoc.csv”.
- Every cell-cell contact recorded in “GuiNeighbor\Embryo_Time_guiNeighbor.txt” has an area recorded in “Embryo_Stat.csv”, while the contacted cells have corresponding segmented regions recorded in “SegCell\Embryo_Time_segCell.nii.gz”.
- Every dividing cell recorded in “DivisionCell\Embryo_Time_division.txt” has no nucleus recorded in “Embryo_CD.csv” but the nuclei of its two daughter cells are recorded.
- Every dividing cell recorded in “DivisionCell\Embryo_Time_division.txt” has a volume recorded in “Embryo_volume.csv” and a surface area recorded in “Embryo_surface.csv”. If its division ends before the terminal edited time point, there are volumes recorded in “Embryo_volume.csv” and surface areas recorded in “Embryo_surface.csv” for its two daughter cells.
- Every dividing cell recorded in “DivisionCell\Embryo_Time_division.txt” is included in “Embryo_lifescycle.csv” and “TPCell\Embryo_Time_cells.txt”, and has a corresponding segmented region recorded in “SegCell\Embryo_Time_segCell.nii.gz”.

## (ii) SUPPLEMENTAL FIGURES

**Fig. S1:**
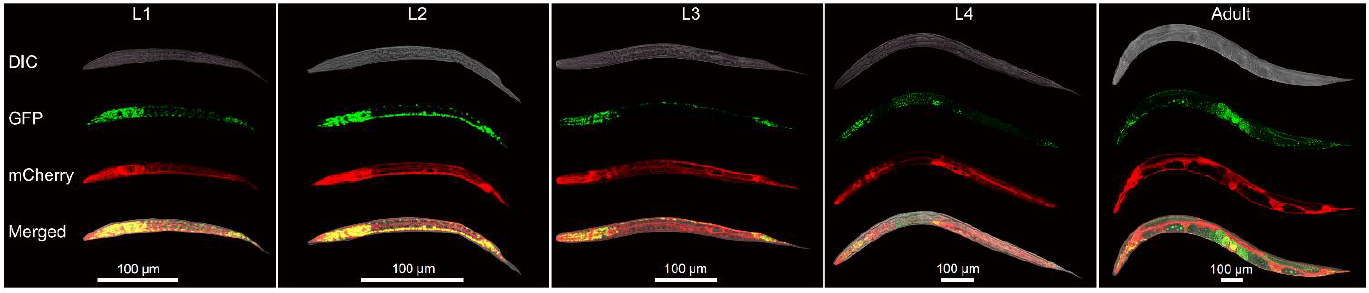
The microscopic imaging on the novel transgenic *C. elegans* strain at its larva (*abbr.*, L) and adult stages. Here, the images for L1, L2, L3, L4, and adult stages are shown from left to right; the images obtained from DIC (Differential Interference Contrast) and through green (Green Fluorescent Protein labeling cell nucleus, *abbr.*, GFP) and red (mCherry labeling cell membrane) fluorescence channels along with their mergence are shown from top to bottom.

**Fig. S2:**
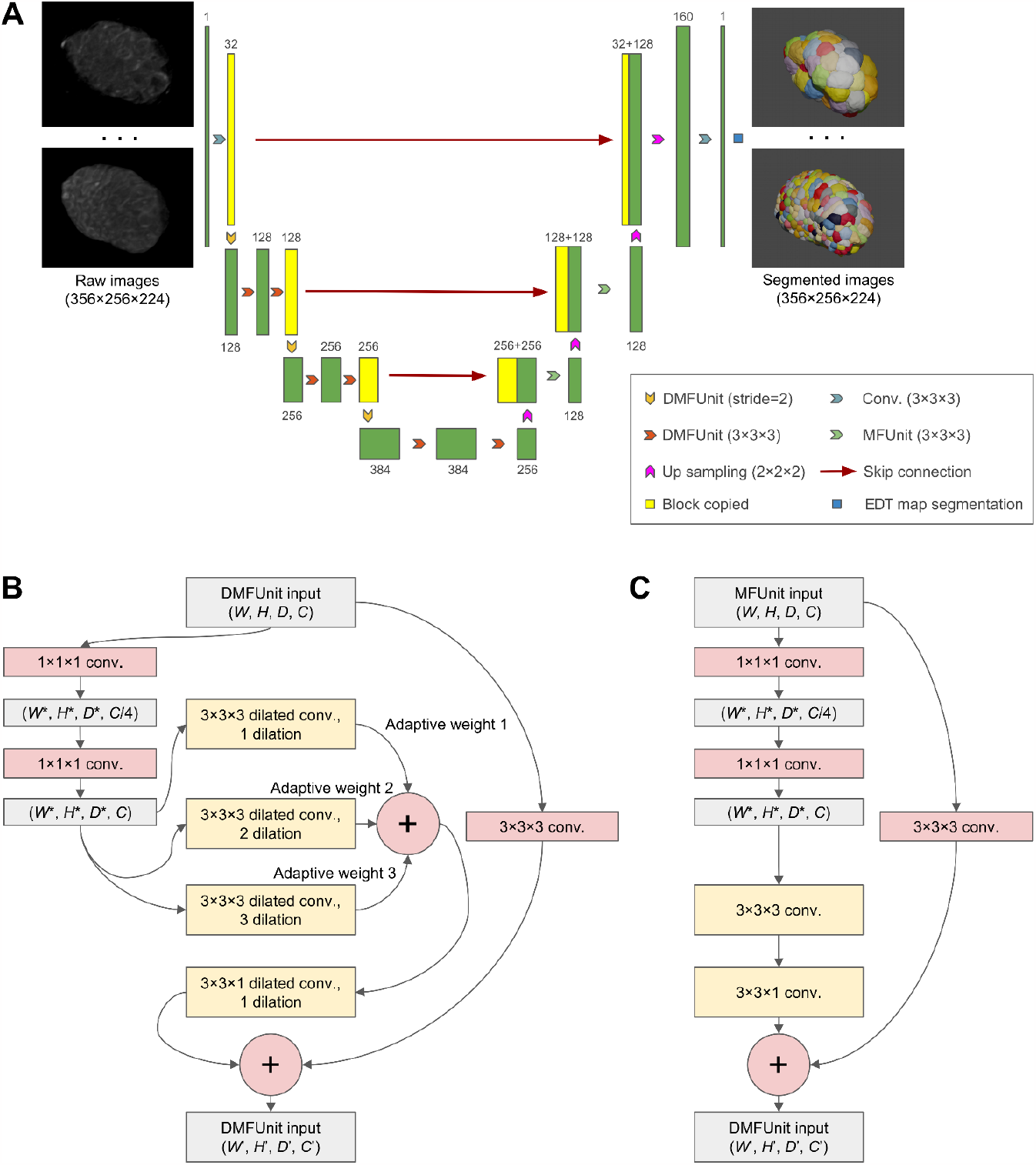
The detailed convolutional neural network structure and data processing flow of *CMap* segmentation. (**A**) The overall U-shape structure of EDT-DMFNet with the channel number of each 3D deep convolutional layer marked. Here, the model conducts semantic segmentation on the *z*-stack of 3D image data and generates segmented 3D cell objects by cell nucleus marker-based watershed algorithm (*i.e.*, EDT map segmentation). (**B**) The detailed inner data flow of DMFUnit of the encoders (the left part of the EDT-DMFNet shown in (**A**)), where dilated convolution cores and multiple fibers are used for exchanging information. (**C**) The detailed inner data flow of MFUnit of the decoders (the right part of the EDT-DMFNet shown in (**A**)).

**Fig. S3:**
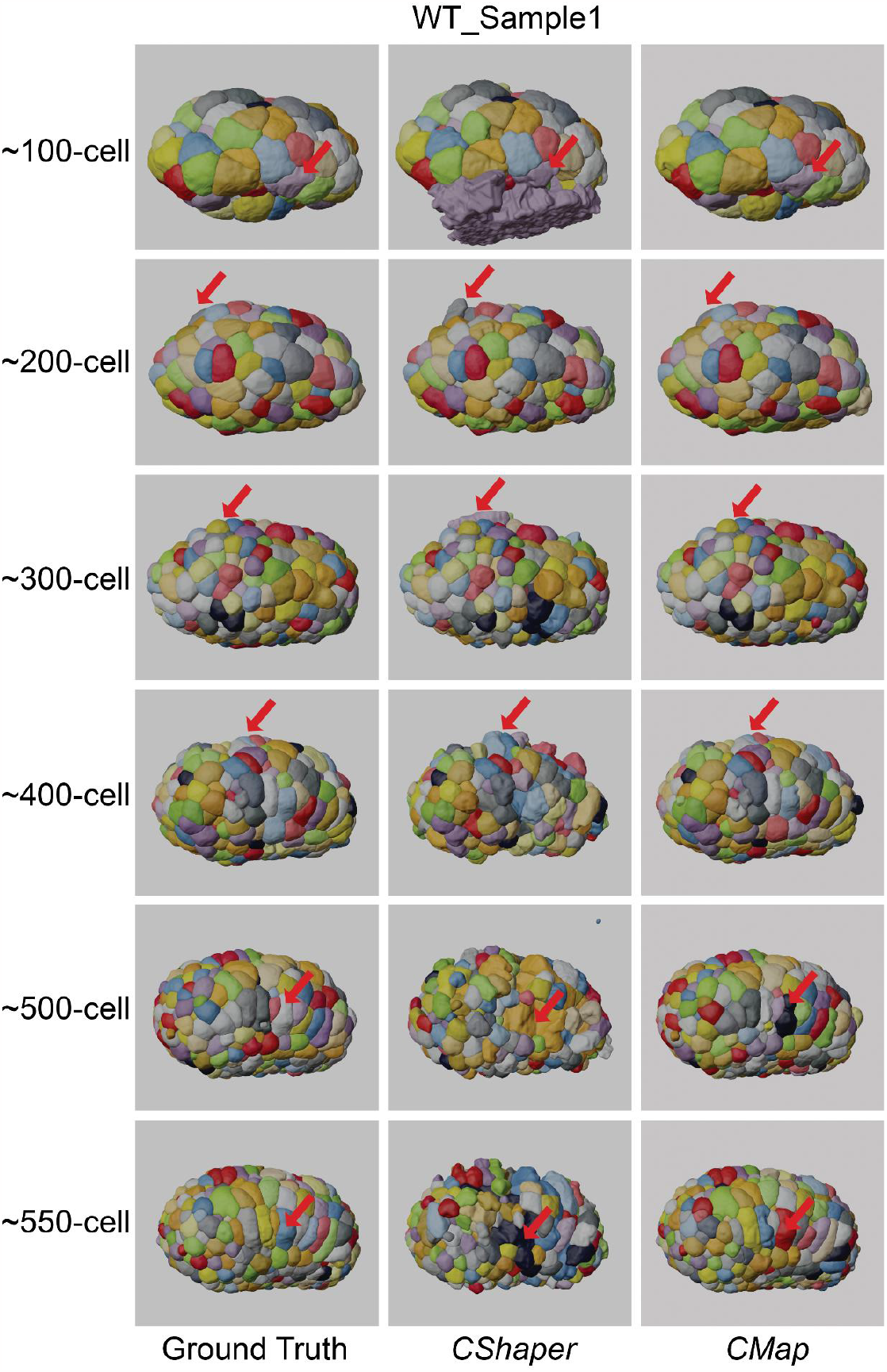
The 3D segmentation comparison between the ground truth (left column), *CShaper* outputs (middle column), and *CMap* outputs (right column), using the embryo sample WT_Sample1. Here, the snapshots at around 100-, 200-, 300-, 400-, 500-, and 550-cell stages are shown from top to bottom (Materials and Methods - Manual annotation); representative segmentation defects in *CShaper* outputs but not in *CMap* outputs are indicated by red arrows.

**Fig. S4:**
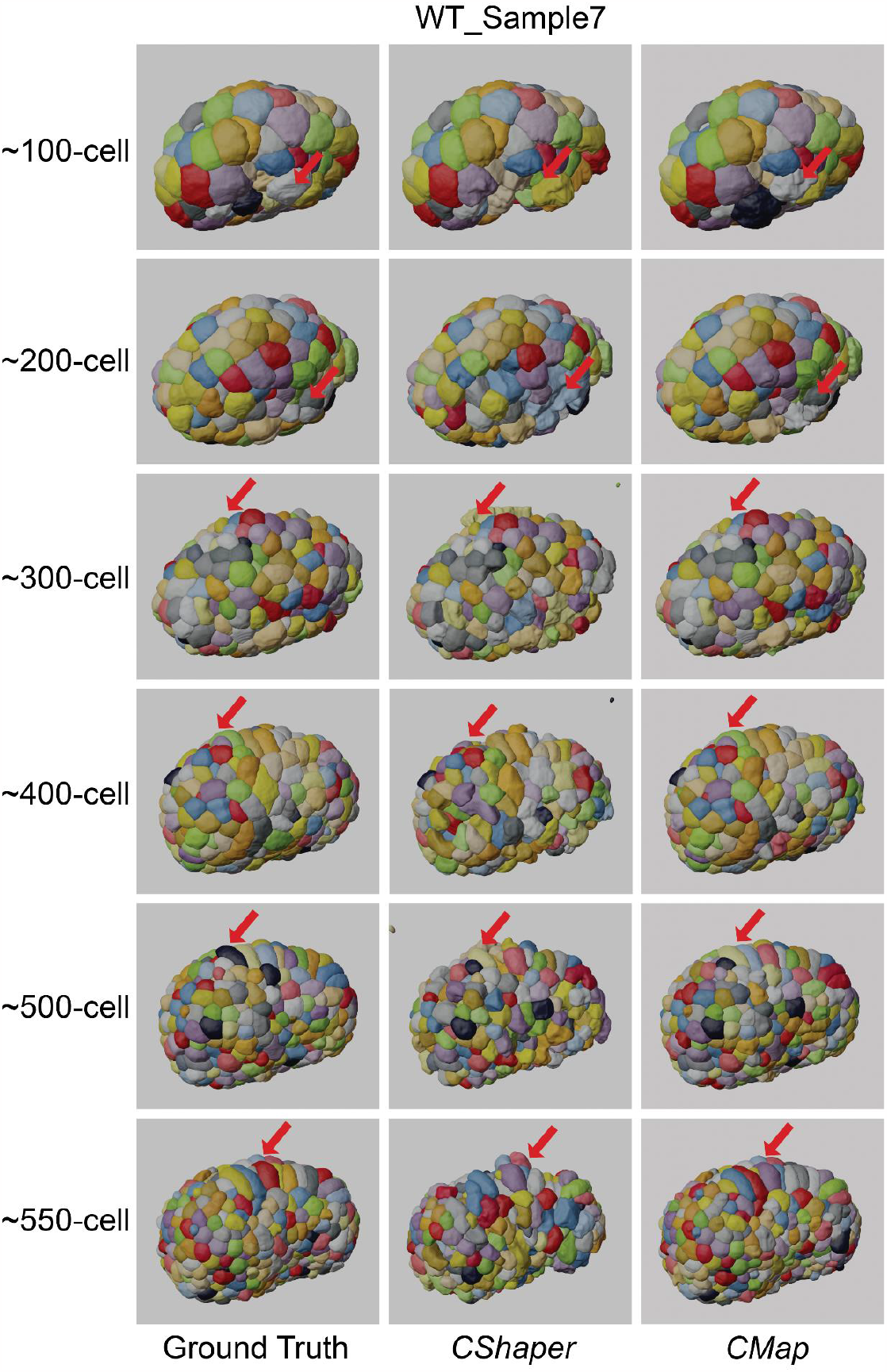
The 3D segmentation comparison between the ground truth (left column), *CShaper* outputs (middle column), and *CMap* outputs (right column), using the embryo sample WT_Sample7. Here, the snapshots at around 100-, 200-, 300-, 400-, 500-, and 550-cell stages are shown from top to bottom (Materials and Methods - Manual annotation); representative segmentation defects in *CShaper* outputs but not in *CMap* outputs are indicated by red arrows.

**Fig. S5:**
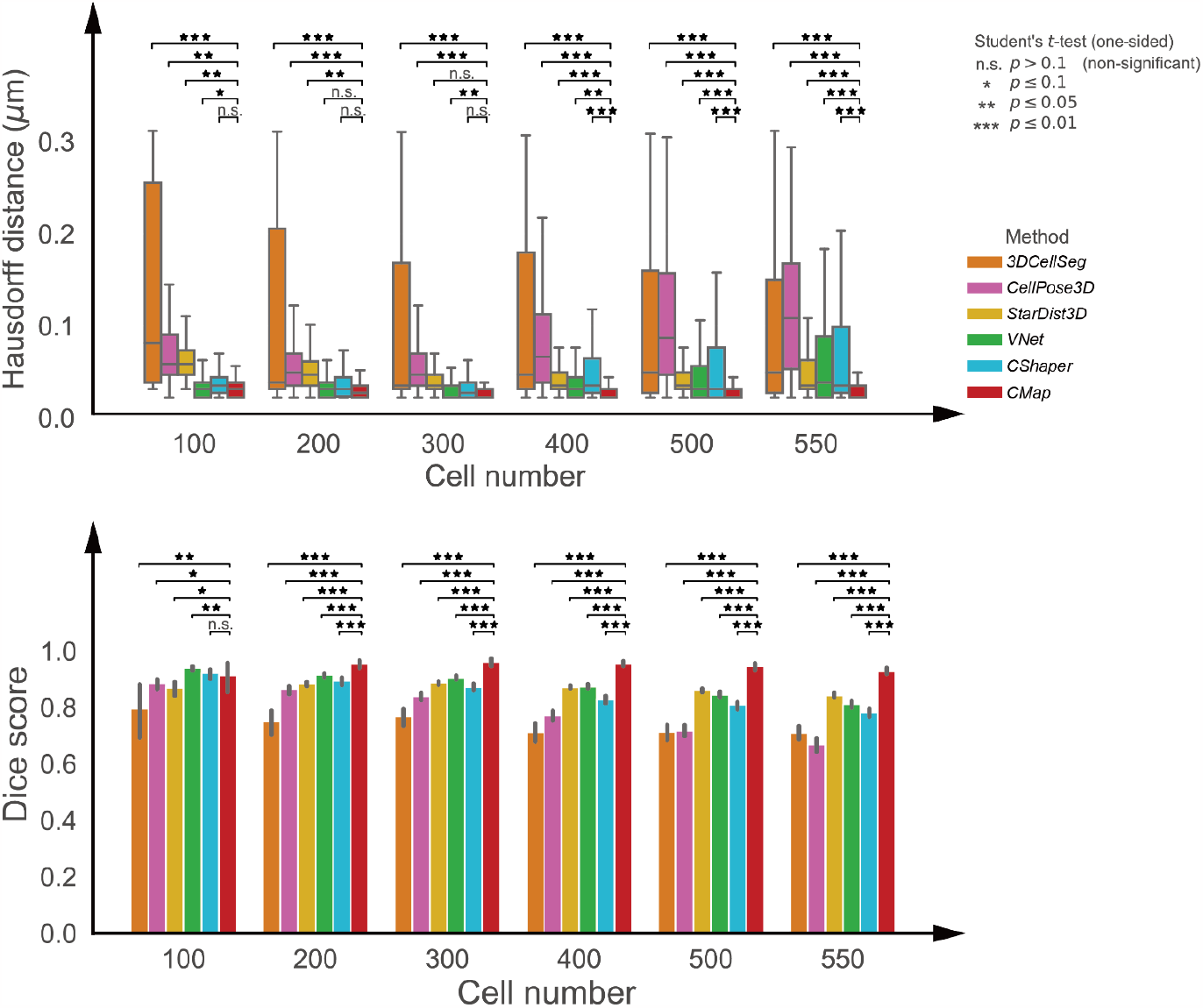
The comparison of Hausdorff distance (upper panel) and Dice score (lower panel) between the state-of-the-art cell segmentation algorithms considered at different developmental stages (represented by cell numbers), using the manually annotated ground truth images as a benchmark.

**Fig. S6:**
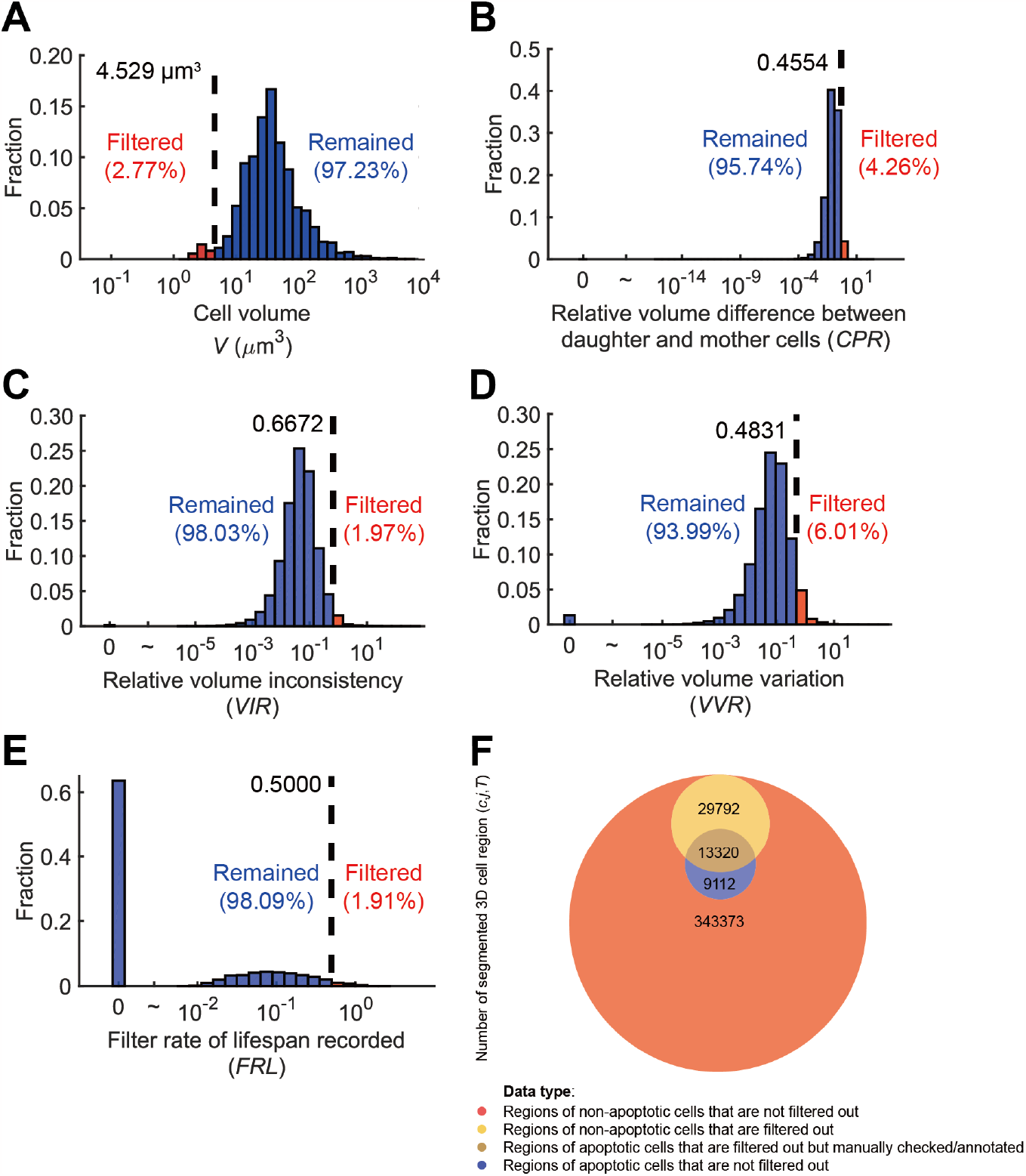
The filtering result of the segmented 3D cell regions from the last time point of the 4-cell stage to the last edited time point. (**A**) The histogram of the volume (*i.e., V*) of the cell regions. (**B**) The histogram of the relative difference in volume between daughter cells and their mother (*i.e., CPR*). (**C**) The histogram of the volume inconsistency of cell *c* at two consecutive time points *T* + 1 and *T* (*i.e., VIR*). (**D**) The histogram of the volume variation of cell *c* at a specific time point *T* (*i.e., VVR*). (**E**) The histogram of the filter rate of lifespan recorded (*i.e., FRL*). (**F**) The pools of segmented 3D cell regions and the ones filtered out or proceeding apoptosis, symbolized by circles with different colors and areas (proportional to number).

**Fig. S7:**
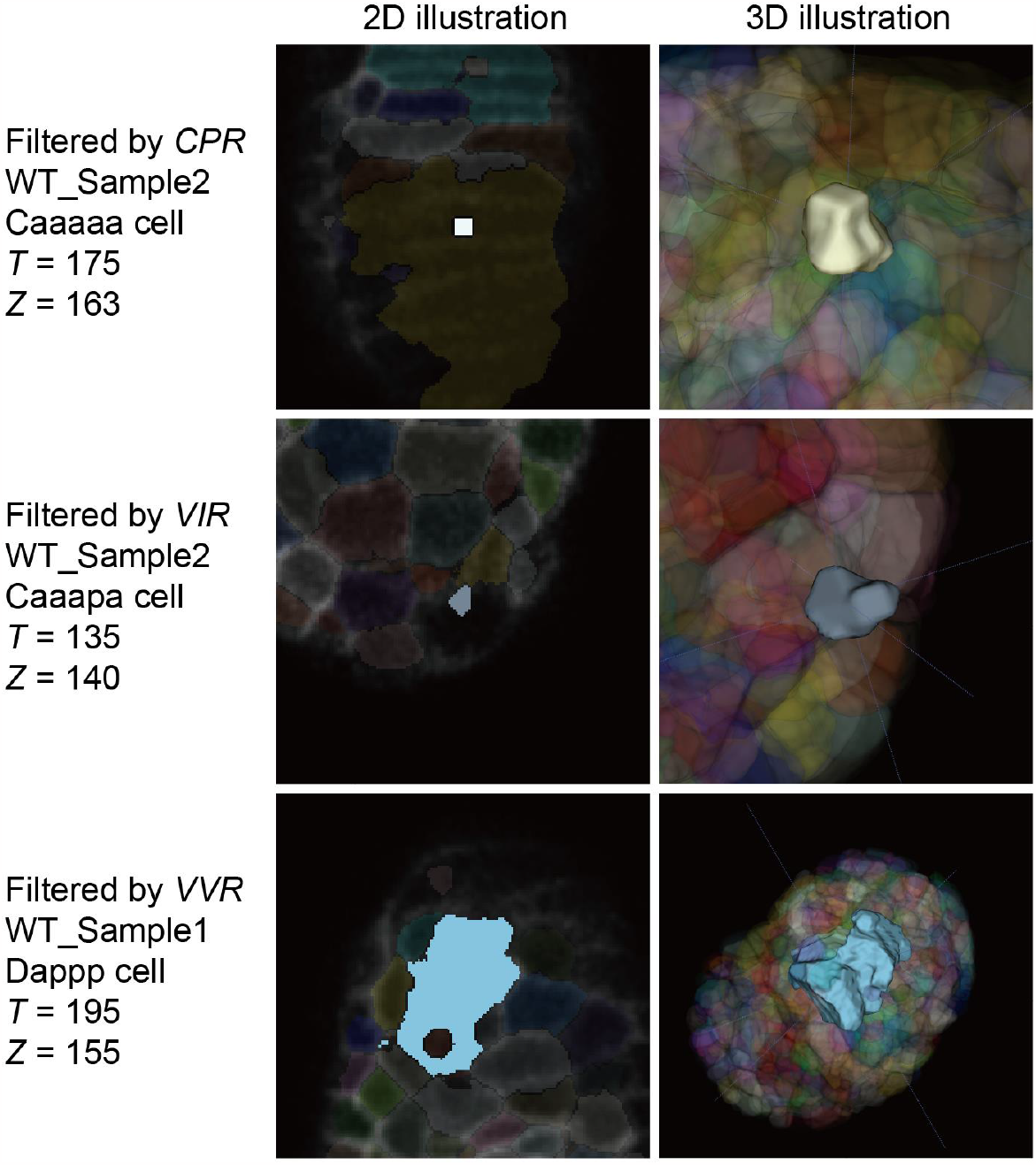
The 2D and 3D snapshots for the exemplary cell regions filtered by *CPR* (upper row), *VIR* (middle row), and *VVR* (lower row). Here, the illustrations are realized with the aid of *ITK-SNAP* [Yushkevich *et al., EMBC*, 2016]; the definition and calculation for *CPR, VIR*, and *VVR* are detailed in Materials and Methods - Data filtering based on abnormal cell volume and cell shape.

**Fig. S8:**
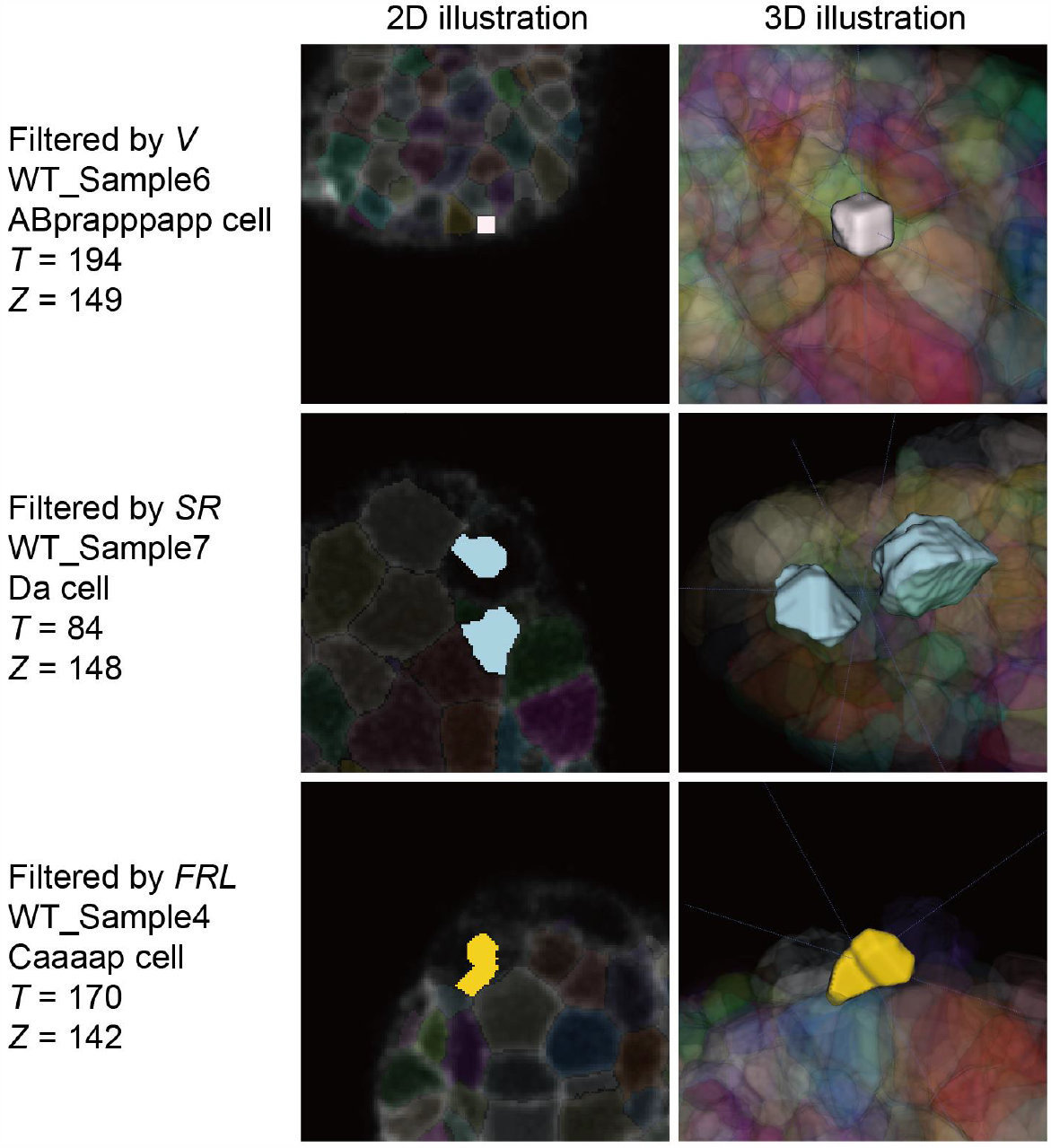
The 2D and 3D snapshots for the exemplary cell regions filtered by *V* (upper row), *SR* (middle row), and *FRL* (lower row). Here, the illustrations are realized with the aid of *ITK-SNAP* [Yushkevich *et al., EMBC*, 2016]; the definition and calculation for *V, SR*, and *FRL* are detailed in Materials and Methods - Data filtering based on abnormal cell volume and cell shape.

**Fig. S9:**
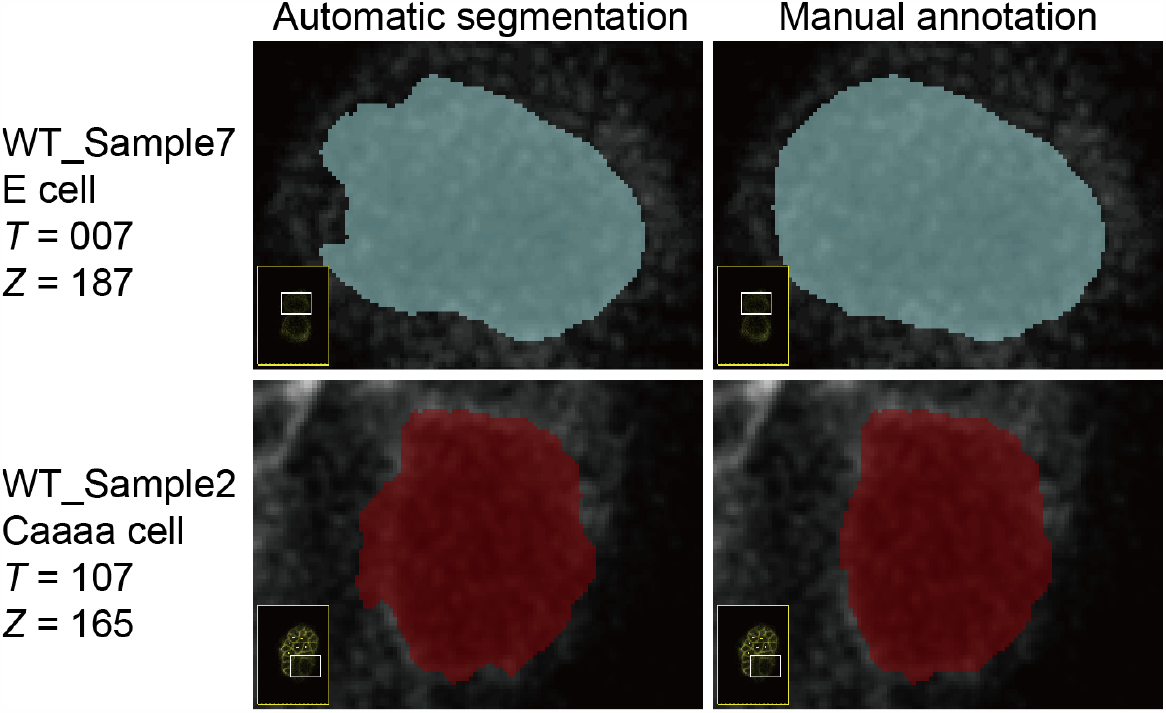
The 2D snapshots for the exemplary non-apoptotic cell regions with defective automatic segmentation (left column) that can be corrected by manual annotation (right column). Here, the illustrations are realized with the aid of *ITK-SNAP* [Yushkevich *et al., EMBC*, 2016].

**Fig. S10:**
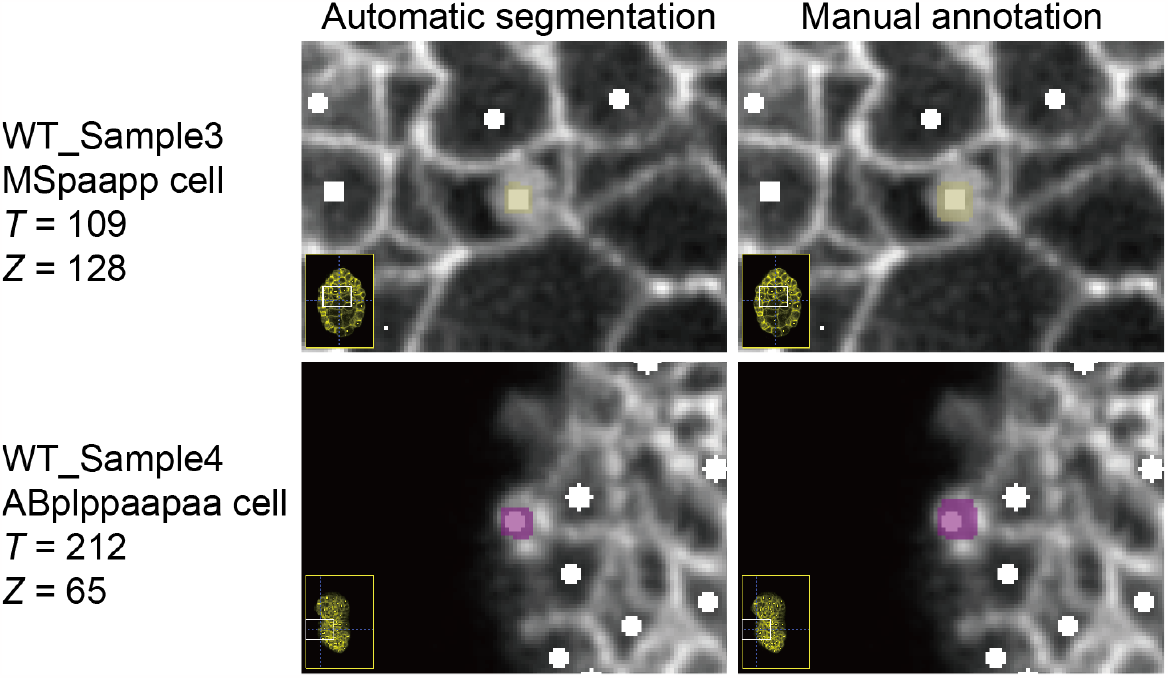
The 2D snapshots for the exemplary apoptotic cell regions with defective automatic segmentation (left column) that can be corrected by manual annotation (right column). Here, the illustrations are realized with the aid of *ITK-SNAP* [Yushkevich *et al., EMBC*, 2016].

**Fig. S11:**
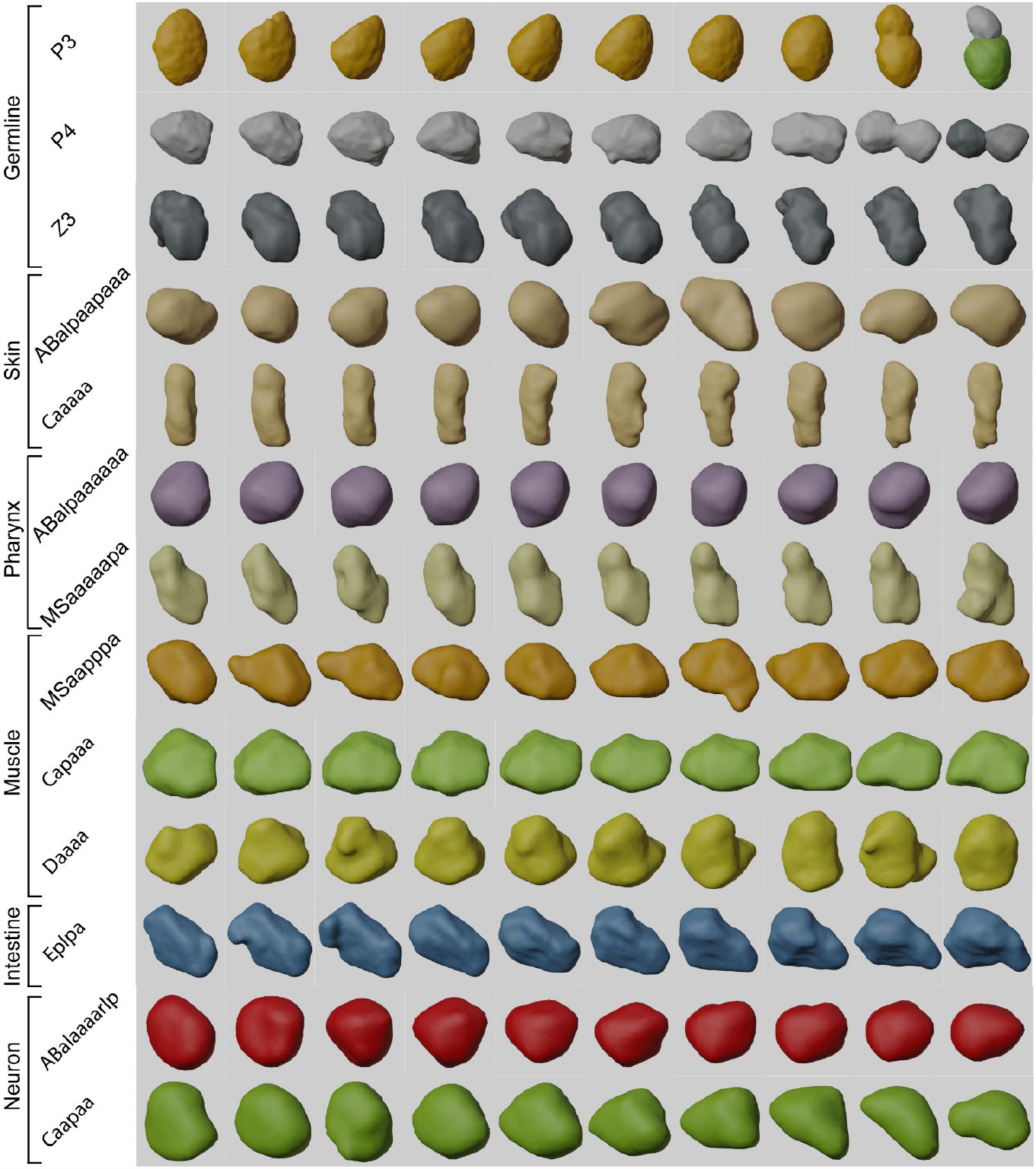
The 3D snapshots for cells from different lineages and with different fates. Here, for the P3 and P4 cells, the last consecutive ten time points before their membrane segregation (included) are shown from left to right; for the other cells, their first consecutive ten time points since the ≥550-cell stage are shown from left to right; the sequential snapshots for a cell are collected from the same embryo sample without data point loss.

**Fig. S12:**
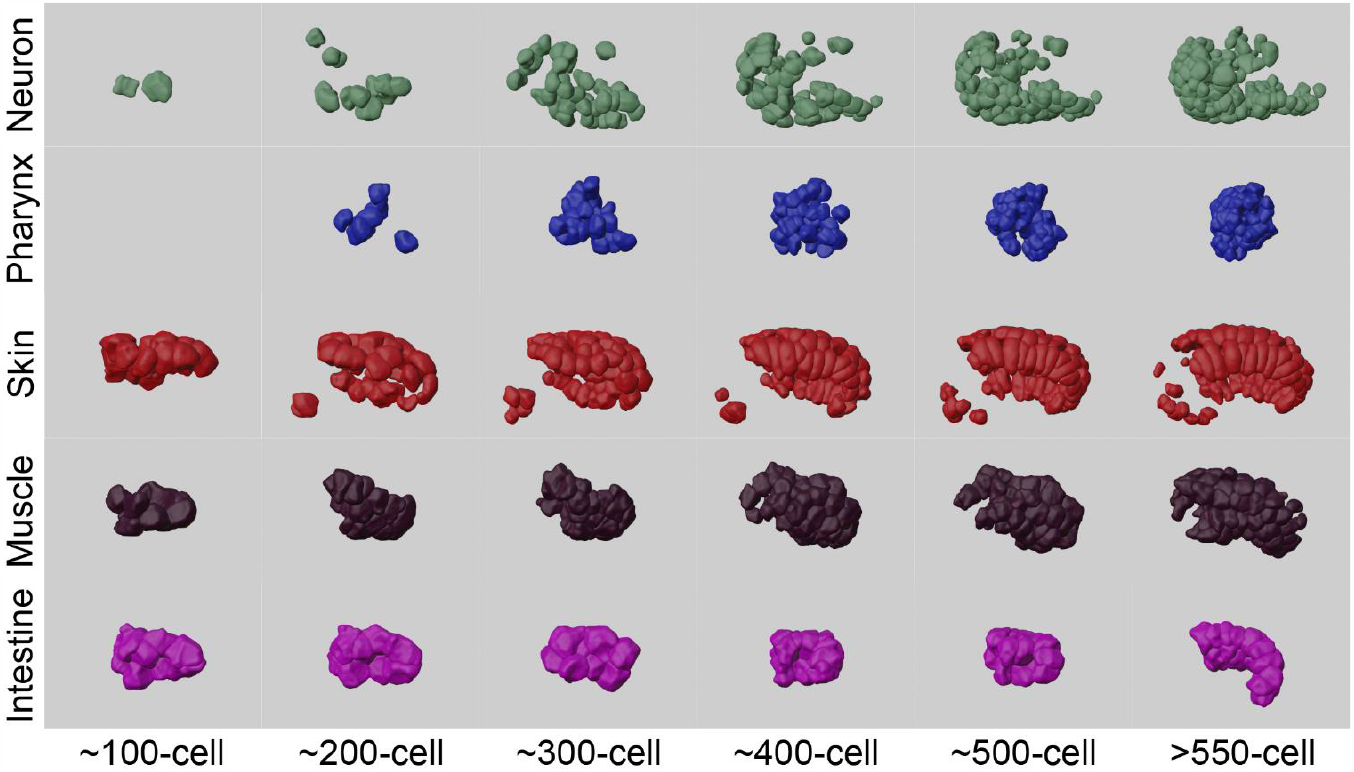
The 3D snapshots for different tissues and organs within the 100±10 -, 200±10 -, 300±10 -, 400±10 -, 500±10 -, and >550-cell stages. Here, the sequential snapshots for a tissue or organ are collected from the same embryo sample; note that the data point loss rate is always lower than 5% for all the snapshots.

**Fig. S13:**
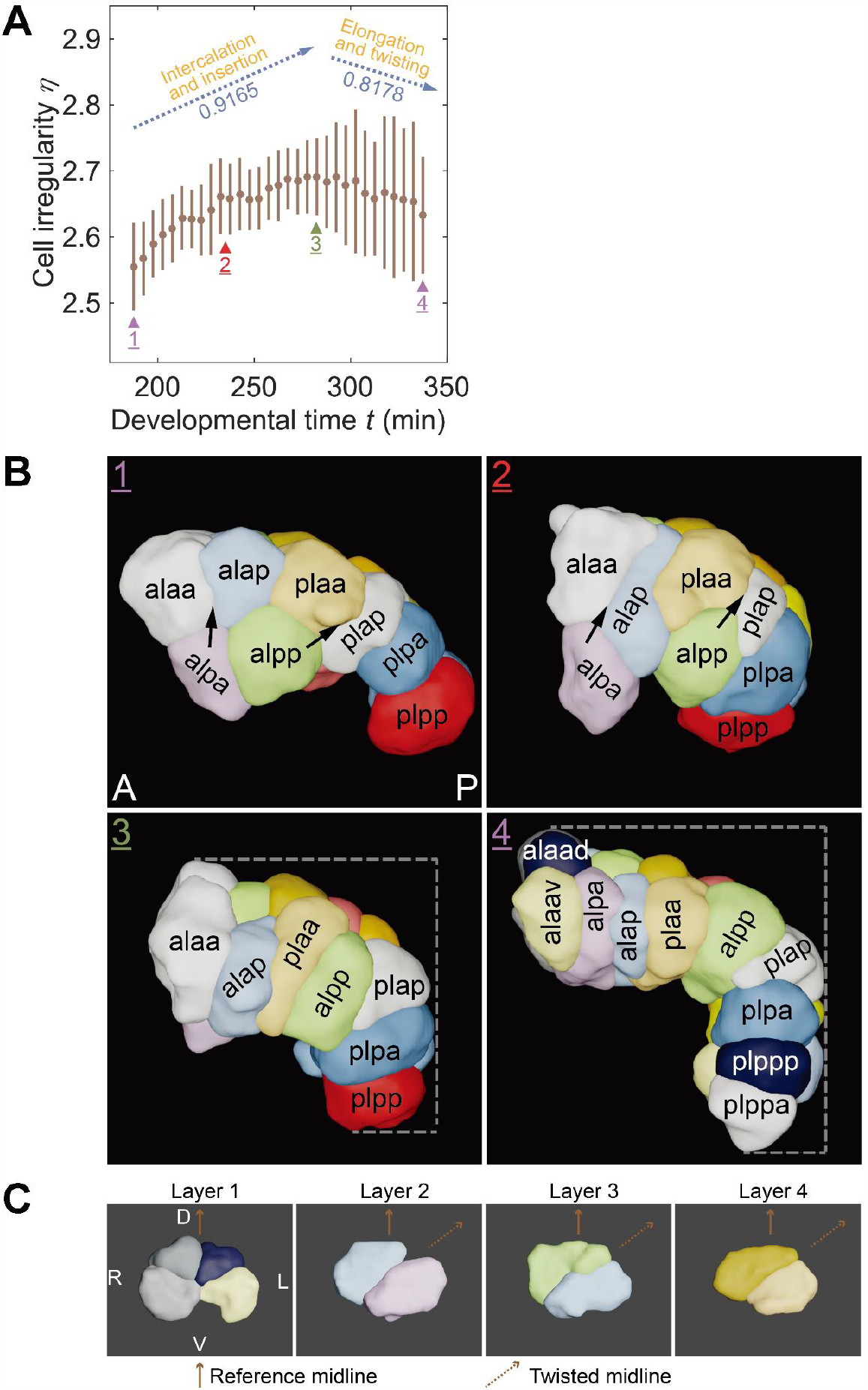
The qualitative and quantitative characterization of the cells participating in intestinal morphogenesis in a *C. elegans* embryo. (**A**) The cell irregularity (η; with the solid circle representing the average and the vertical line representing the standard deviation within a constant interval) dynamic of the cells participating in intestinal morphogenesis over developmental time (*t*; with the last time point of the 4-cell stage as the time zero), in accordance with Figure 3C and same as Figure 3F. Here, all the E cells are taken into account; the global maximum of the η − *t* curve is indicated by a green triangle where the correlation coefficient of the monotonic partial curves it partitions are labeled at the top; note that Moment 1 is the first time point when the E16 cells co-exist and Moment 4 is the last time point when the E20 cells co-exist; Moment 3 is the time point when the cell irregularity reaches its global maximum, then Moment 2 is the middle time point between Moment 1 and Moment 3. (**B**) The 3D snapshots for the cells participating in intestinal morphogenesis at the 4 moments labeled in (**A**). Here, Moment 1 and Moment 2 show the intercalation and insertion of particular E cells (*i.e.*, the Ealpa, Earpa, Ealpp, Earpp cells), while Moment 3 and Moment 4 show the elongation and twisting of the entire intestine. (**C**) The twisting of the first four cell layers, highlighted by the reference and twisted midlines; Here, consistent with Moment 4 in (**B**), Layer 1 includes the Ealaad, Ealaav, Earaad, and Earaav cells; Layer 2 includes the Ealpa and Earpa cells; Layer 3 includes the Ealap and Earap cells; Layer 4 includes the Eplaa and Epraa cells.

**Fig. S14:**
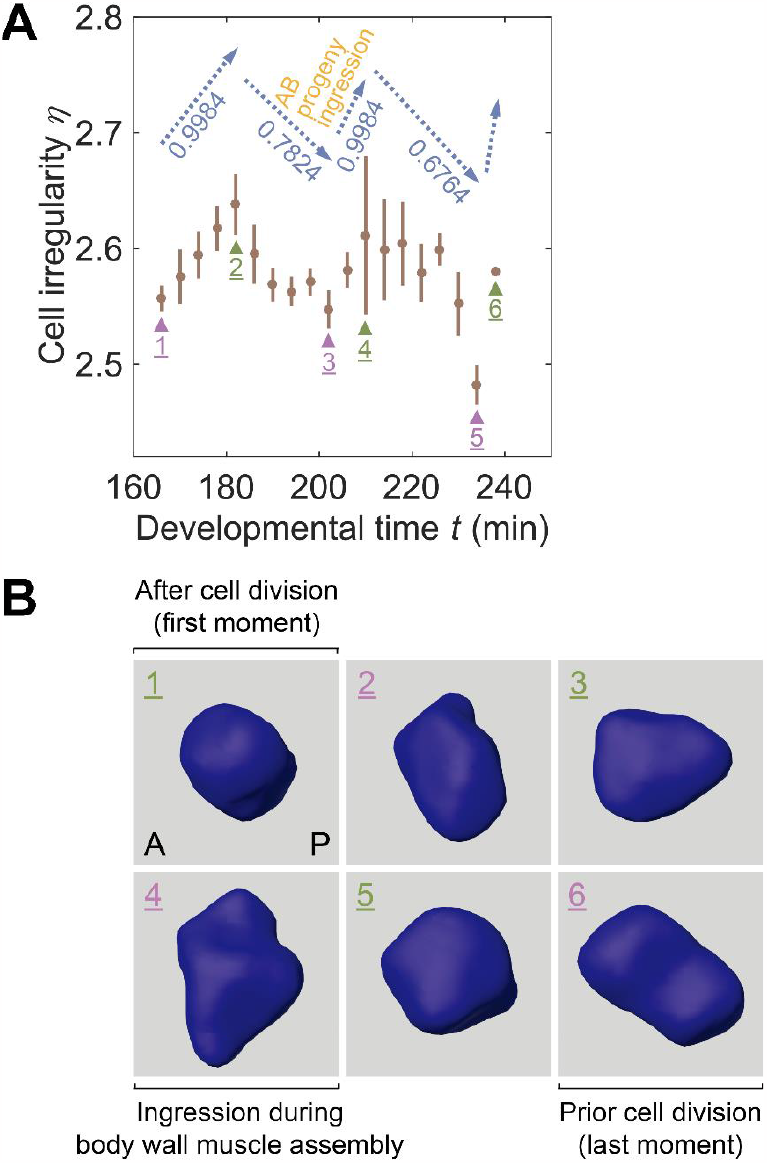
The qualitative and quantitative characterization of the AB progeny (*i.e.*, the ABprpppppaa cell) participating in body wall muscle assembly in a *C. elegans* embryo. (**A**) The cell irregularity (η; with the solid circle representing the average and the vertical line representing the standard deviation within a constant interval) dynamic of the AB progeny (*i.e.*, the ABprpppppaa cell) participating in body wall muscle assembly over developmental time (*t*; with the last time point of the 4-cell stage as the time zero), in accordance with Figure 3D. Here, only the data of the *C. elegans* wild-type embryo sample WT_Sample1 is taken into account; the local maximums and minimums of the η − *t* curve are indicated by green and purple triangles respectively, where the correlation coefficient of the monotonic partial curves they partition are labeled at the top. (**B**) The 3D snapshots for the AB progeny (*i.e.*, the ABprpppppaa cell) participating in body wall muscle assembly at the six moments labeled in (**A**).

**Fig. S15:**
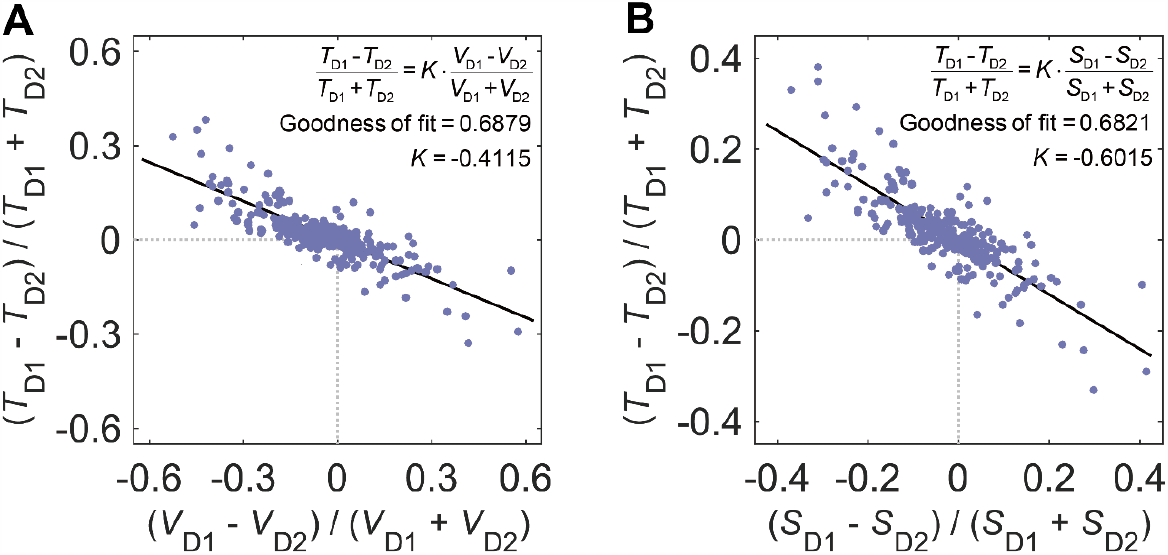
The anti-correlation between cell size asymmetry and cell cycle length asymmetry. (**A**) The anti-correlation between cell volume asymmetry (*i.e.*,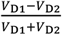) and cell cycle length asymmetry (*i.e.*,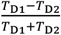). Here, the result of proportional fitting between 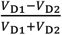 and 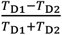is shown with a solid line, with the proportional coefficient (*i.e., K*) and goodness of fit (*i.e., G*) listed in the top right corner; a total of 285 cells recorded with a complete lifespan for their daughter cells and themselves are taken into account (Table S9). (**B**) The anti-correlation between cell volume asymmetry (*i.e.*,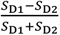) and cell cycle length asymmetry (*i.e.*, 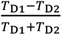). Here, the result of proportional fitting between 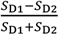 and 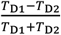 is shown with a solid line, with the proportional coefficient (*i.e., K*) and goodness of fit (*i.e., G*) listed in the top right corner; a total of 285 cells recorded with a complete lifespan for their daughter cells and themselves are taken into account (Table S9).

**Fig. S16:**
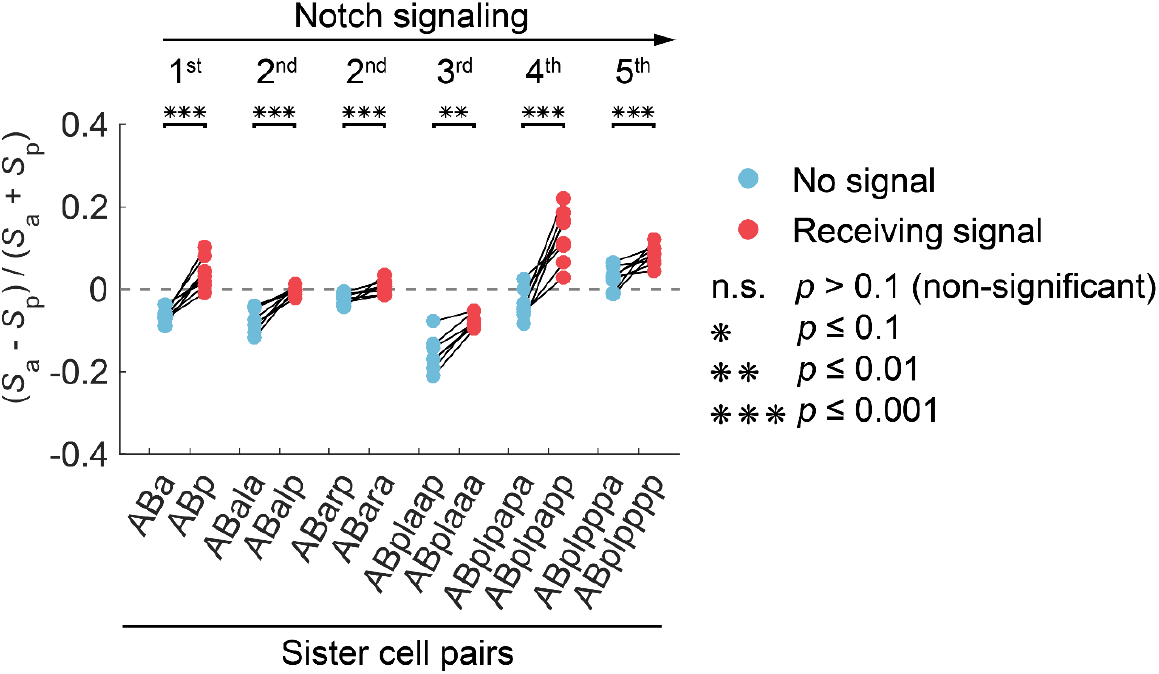
The cell surface area asymmetry between daughter cells (*i.e.*,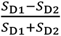) in the six sister cell pairs differentiated through Notch signaling. Here, for each of the six sister cell pairs, the cell with no signaling is painted in blue while the cell receiving signaling is painted in red; the data from all eight wild-type embryo samples are presented, where the two data points of a sister cell pair from the same embryo samples are connected with a solid black line; the statistical significance is obtained by the one-sided Wilcoxon rank-sum test (non-significant, *abbr.*, n.s., *p* > 0.1; *, *p* ≤ 0.1; **, *p* ≤ 0.01; ***, *p* ≤ 0.001) and is listed on the right.

**Fig. S17:**
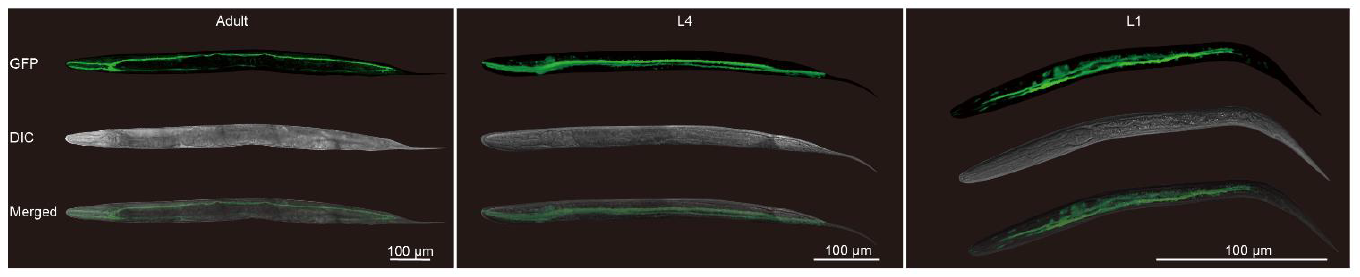
The H-shaped kidney cell fluorescently labeled by GFP at the adult, L4, and L1 stages [Zhao *et al., J. Biol. Chem.*, 2005], illustrated with the images from confocal microscopy (upper row) and differential interference contrast microscopy (middle row) and their mergence (lower row). Here, a scale bar of 100 μm is plotted in the bottom right corner of the subfigure for each stage.

**Fig. S18:**
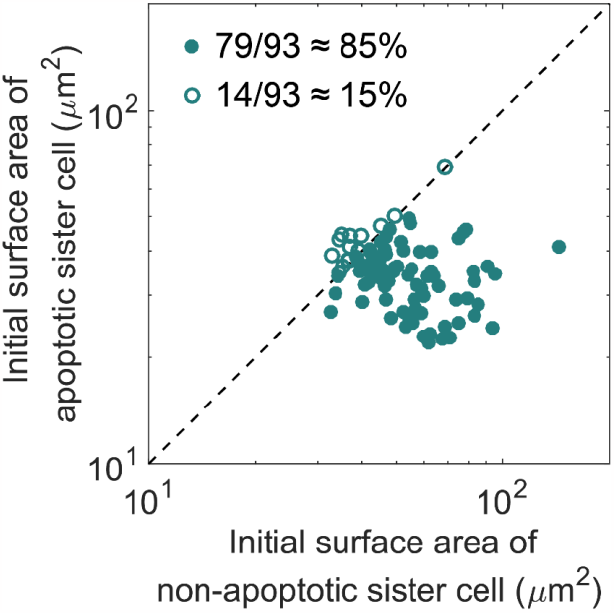
The smaller initial surface area of an apoptotic sister cell (vertical coordinate) compared to the one of its non-apoptotic sister cell (horizontal coordinate). Here, the data of a total of 93 properly-recorded sister cell pairs are presented, among which 79 have a relatively smaller initial volume for their apoptotic sister cells on average.

**Fig. S19:**
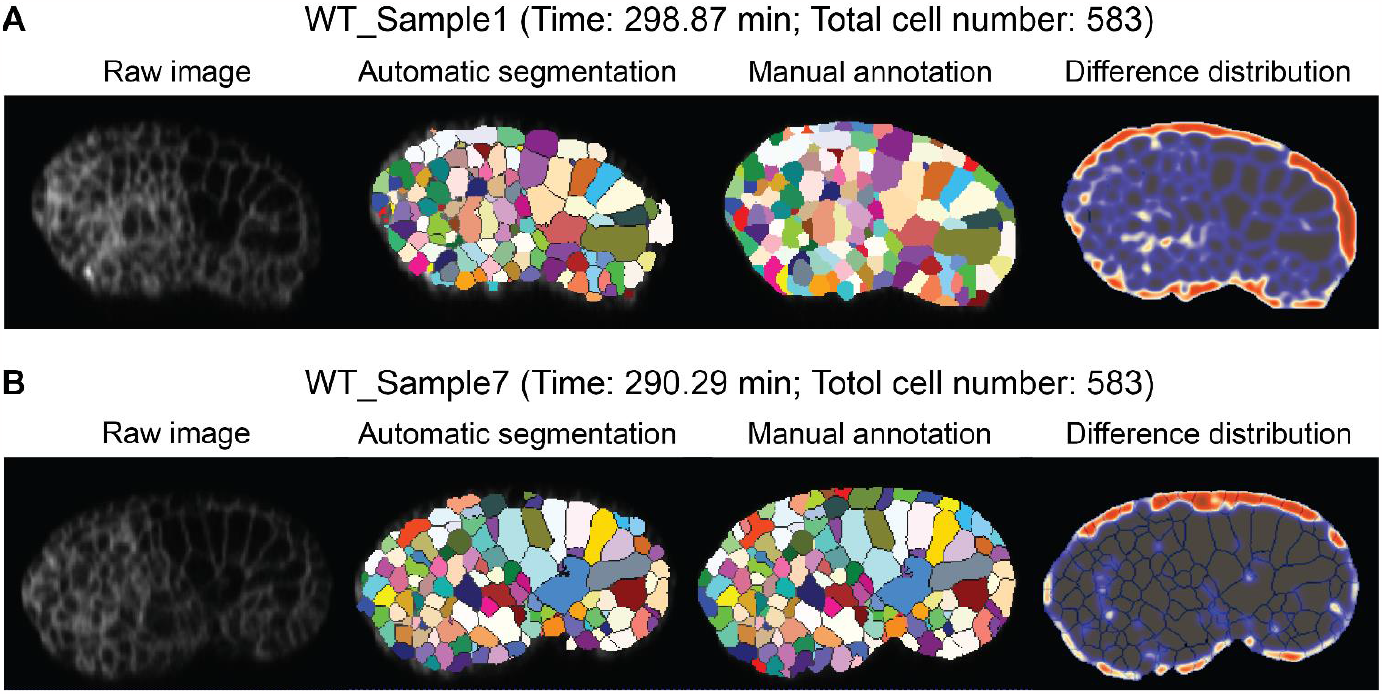
The segmentation accuracy of *CMap* beyond the 550-cell stage. (**A**) The embryo sample WT_Sample1 with the absolute imaging time and total cell number listed at the top. Here, the raw image is placed in the 1^st^ column, while its automatic segmentation and manual annotation are shown in the 2^nd^ and 3^rd^ columns followed by the distribution of their difference (highlighted in red) in the 4^th^ column. (**B**) The embryo sample WT_Sample7 with the absolute imaging time and total cell number listed at the top. Here, the raw image is placed in the 1^st^ column, while its automatic segmentation and manual annotation are shown in the 2^nd^ and 3^rd^ columns followed by the distribution of their difference (highlighted in red) in the 4^th^ column.

**Fig. S20:**
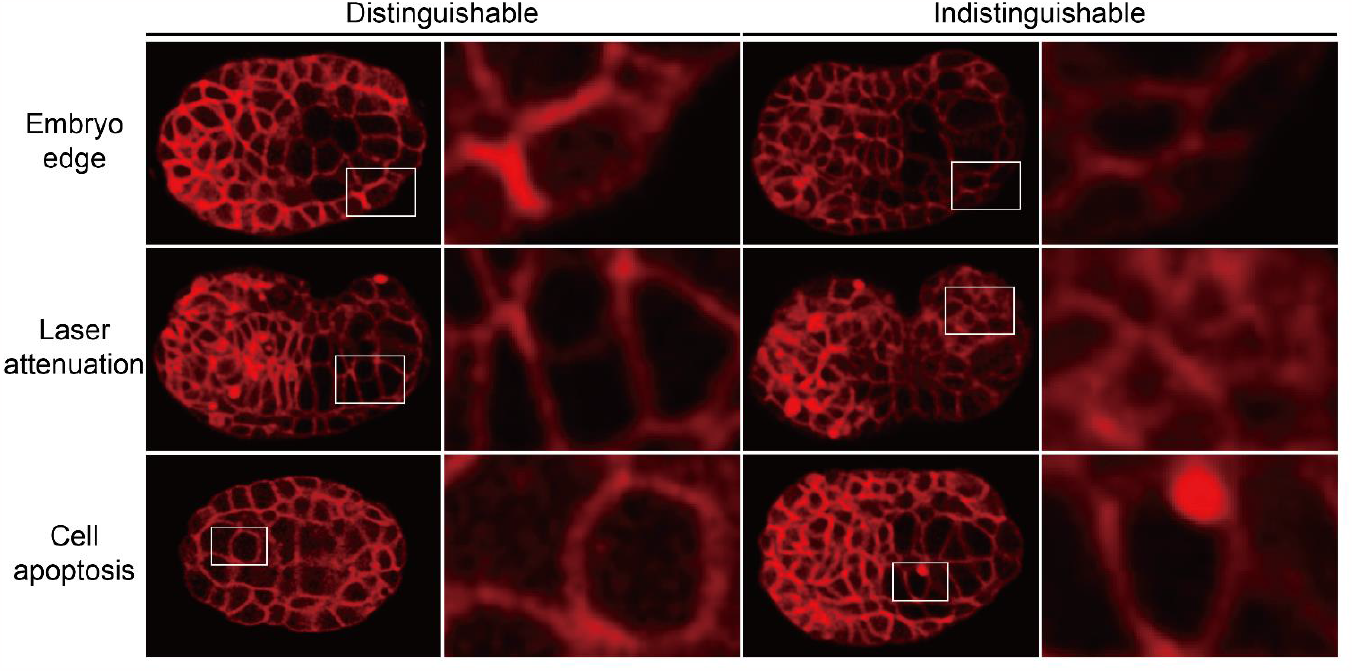
The blurred region exemplified by the fluorescence images captured in the condition with amplified laser intensity for one shot (left panel; distinguishable) and in the actual condition (right; indistinguishable). Here, for each row with a specific interference factor and in each condition, the comparable region is indicated by a white rectangle in the overall image on the left and it’s zoomed in on the right.

**Fig. S21:**
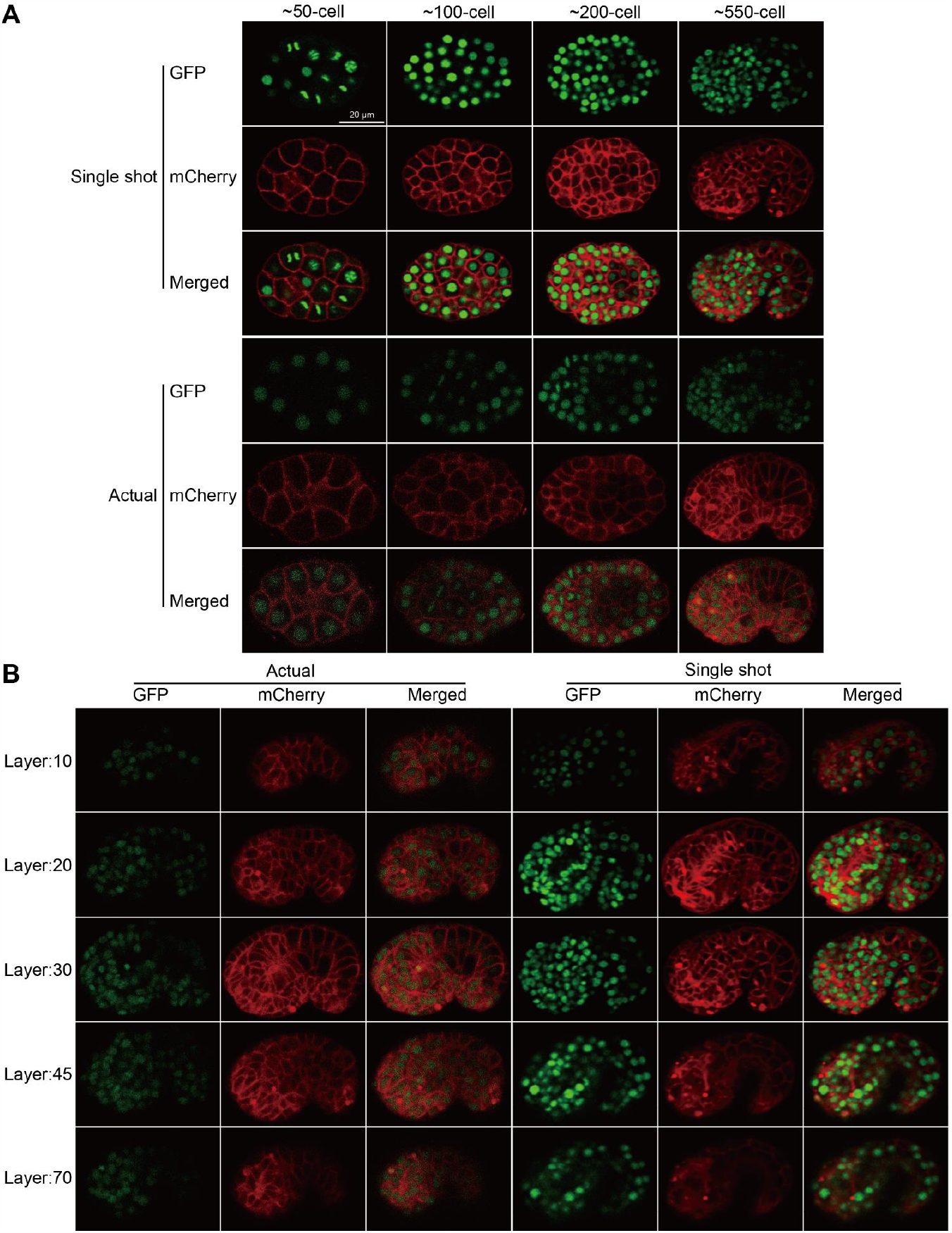
The fluorescence images captured in the actual condition and in the one with amplified laser intensity for one shot (with a scale bar of 20 μm plotted in the top left corner). (**A**) The fluorescence images near the middle layer at the roughly 50, 100, 200, and 550-cell stages. (**B**) The fluorescence images in the 10^th^, 20^th^, 30^th^, 45^th^, and 70^th^ layers at the roughly 550-cell stage.

## (iii) CAPTIONS FOR SUPPLEMENTAL TABLES

**Table S1: The computational time cost of *CMap*for segmenting a *C. elegans*embryo from the ≤4- to >550-cell stages.**

**Table S2: The variation between automatic segmentation and ground truth.**

**Table S3: The experimental information of embryo samples.**

**Table S4: The cell list in different standards and conditions.**

**Table S5: The comparison between *CShaper*and *CMap*datasets.**

**Table S6: The lost and annotated cell list.**

**Table S7: The data point loss rate regarding time.**

**Table S8: The data point loss rate regarding cell identity, cell lineage, and cell fate.**

**Table S9: The asymmetry of cell size and cell cycle length.**

**Table S10: The geometric estimation for compressed eggshells.**

**Table S11: The cell volume asymmetry change between uncompressed and compressed embryos.**

**Table S12: The initial size of non-apoptotic and apoptotic sister cell pairs.**

**Table S13: The format of multidimensional data outputted from *CMap***.

## (iv) CAPTIONS FOR SUPPLEMENTAL MOVIES

**Movie S1: The 3D morphology reconstruction of all cells in a *C. elegans* wild-type embryo sample, shown by the time points lasting from the <4- to >550-cell stages, with cell-wise color scheme.** Here, the upper morphologies from left to right are observed along the *z, y*, and *x* axes, respectively; the lower morphologies from left to right are the crosssection in the middle layer of the upper morphologies.

**Movie S2: The 3D morphology reconstruction of all cells in a *C. elegans* wild-type embryo sample, shown by the time points lasting from the <4- to >550-cell stages, using a lineage-wise color scheme.** Here, the cell lineages include the AB, MS, E (*incl.*, the E cells and EMS cell), C, D, and P (*incl.*, the P1, P2, P3, P4, and Z2 and Z3 cells), according to previous literature [Sulston *et al., Dev. Biol.*, 1983].

**Movie S3: The 3D morphology reconstruction of all cells in a *C. elegans* wild-type embryo sample, shown by the time points lasting from the <4- to >550-cell stages, using a fate-wise color scheme.** Here, the cell fates include the neuron, pharynx, intestine, germline, muscle, skin, death, unspecified, and others, according to previous literature [Sulston *et al., Dev. Biol.*, 1983; Li *et al., Cell Rep.*, 2019].

**Movie S4: The 3D morphology reconstruction of the cells with five different fates (*incl.*, the neuron, pharynx, intestine, muscle, skin), shown by a time point at the >550-cell stage with a data point loss rate lower than 5%, using a cell-wise color scheme.** Here, the body axes of the embryo are shown at the bottom.

**Movie S5: The 3D morphology reconstruction of the intestinal cells (*i.e.*, the E cells) and germline cells (*i.e.*, the Z2 and Z3 cells) in a *C. elegans* wild-type embryo sample, shown by a time point at the >550-cell stage and without data point loss, using a cell-wise color scheme.** Here, the left panel contains only the intestinal cells; the right panel contains both the intestinal cells (semi-transparent) and germline cells (non-transparent).

**Movie S6: The 3D morphology reconstruction for the gastrulation process in a *C. elegans* wild-type embryo sample, shown by the time points lasting from the 12- to >550-cell stages, using a lineage-wise color scheme.** Here, the cell lineages include the AB, MS, E, C, D, P (*incl.*, the P2, P3, P4 cells) lineages, and the Z2 and Z3 cells, according to previous literature [Sulston *et al., Dev. Biol.*, 1983].

**Movie S7: The 3D morphology reconstruction for the dorsal interdigitation in a *C. elegans* wild-type embryo sample, shown by the time points lasting for >1 hour, using a position-wise color scheme.**

**Movie S8: The 3D morphology reconstruction for the intestinal morphogenesis in a *C. elegans* wild-type embryo sample, shown by the time points from the first time point when the E16 cells co-exist to the terminal edited time point at the >550-cell stage, using a cell-wise color scheme.**

**Movie S9: The 3D morphology reconstruction for the body wall muscle assembly in a *C. elegans* wild-type embryo sample, shown by the time points lasting from the <4- to >550-cell stages, using a lineage-wise color scheme where the ABprpppppaa cell is painted in blue and the ABprpppppap cell is not illustrated.**

**Movie S10: The 3D morphology reconstruction for the body wall muscle assembly in a *C. elegans* wild-type embryo sample, shown by the time points lasting from the <4- to >550-cell stages, using a lineage-wise color scheme where the ABprpppppaa cell is painted in blue and the ABprpppppap cell is painted in gray.**

**Movie S11: The 3D morphology reconstruction of the kidney cell and its ancestors as well as their sister cells in embryo sample WT_Sample1, shown by the time points lasting from the <4- to >550-cell stages, where the apoptotic ABplpappap cell is engulfed by the ABarapappp cell.**

**Movie S12: The 3D morphology reconstruction of the kidney cell and its ancestors as well as their sister cells in embryo sample WT_Sample5, shown by the time points lasting from the <4- to >550-cell stages, where the apoptotic ABplpappap cell is engulfed by the ABplpappaa cell.**

## (vi) AUTHOR CONTRIBUTIONS

H.Y, C.T., and Z.Z. conceived and supervised this project; Y.M., M.-K.W., and L.-Y.C. cultured, imaged, and curated the *C. elegans*; Z.L. and J.C. devised the *CMap* algorithm, performed cell segmentation, and extracted morphological features; G.G., Z.L., and J.C. arranged the manual annotation for ground truth, performed data quality evaluation and control, and organized the datasets; G.G. carried out the quantitative and statistical analyses; Z.L. and G.G. generated the 3D illustration; G.G., Z.L., Y.M., J.C., M.-K.W., and L.-Y.C. designed and improved the *CVE* software; G.G., Z.L., and Y.M. wrote the manuscript; H.Y, C.T., and Z.Z. revised the manuscript.

## Notes

### Competing Interest Statement

The authors have declared no competing interest.

## REFERENCES AND NOTES

1. J. Cao et al., Establishment of a morphological atlas of the Caenorhabditis elegans embryo using deeplearning-based 4D segmentation. Nat. Commun. 11, 6254 (2020). doi: 10.1038/s41467-020-19863-x; pmid: 33288755

2. C. Luxenburg, R. Zaidel-Bar, From cell shape to cell fate via the cytoskeleton — Insights from the epidermis. Exp. Cell Res. 378, 232–237 (2019). doi: 10.1016/j.yexcr.2019.03.016; pmid: 30872138

3. C.Z. Eddy, H. Raposo, A. Manchanda, R. Wong, F. Li, B. Sun, Morphodynamics facilitate cancer cells to navigate 3D extracellular matrix. Sci. Rep. 11, 20434 (2021). doi: 10.1038/s41598-021-99902-9; pmid: 34650167

4. P. Gómez -Gálvez et al., Scutoids are a geometrical solution to three-dimensional packing of epithelia. Nat. Commun. 9, 2960 (2018). doi: 10.1038/s41467-018-05376-1; pmid: 30054479

5. D. Witvliet et al., Connectomes across development reveal principles of brain maturation. Nature 596, 257–261 (2021). doi: 10.1038/s41586-021-03778-8; pmid: 34349261

6. L. Guignard et al., Contact area-dependent cell communication and the morphological invariance of ascidian embryogenesis. Science 369, 158 (2020). doi: 10.1126/science.aar5663; pmid: 32646972

7. R. Fickentscher, P. Struntz, M. Weiss, Setting the clock for fail-safe early embryogenesis. Phys. Rev. Lett. 117, 188101 (2016). doi: 10.1103/PhysRevLett.117.188101; pmid: 27835015

8. J. Priess, Notch signaling in the C. elegans embryo. WormBook (2005). doi: 10.1895/wormbook.1.4.1; pmid: 18050407

9. G. Guan, Z. Zhao, C. Tang, Delineating the mechanisms and design principles of Caenorhabditis elegans embryogenesis using in toto high-resolution imaging data and computational modeling. Comput. Struct. Biotechnol. J. 20, 5500–5515 (2022). doi: 10.1016/j.csbj.2022.08.024; pmid: 36284714

10. Y. Azuma, S. Onami, Biologically constrained optimization based cell membrane segmentation in C. elegans embryos. BMC Bioinform. 18, 307 (2017). doi: 10.1186/s12859-017-1717-6; pmid: 28629355

11. J. Stegmaier et al., Real-time three-dimensional cell segmentation in large-scale microscopy data of developing embryos. Dev. Cell 36, 225–240 (2016). doi: 10.1016/j.devcel.2015.12.028; pmid: 26812020

12. J.E. Sulston, E. Schierenberg, J.G. White, J.N. Thomson, The embryonic cell lineage of the nematode Caenorhabditis elegans. Dev. Biol. 100, 64–119 (1983). doi: 10.1016/0012-1606(83)90201-4; pmid: 6684600

13. J. Nance, J.R. Priess, Cell polarity and gastrulation in C. elegans. Development 129, 387–397 (2002). doi: 10.1242/dev.129.2.387; pmid: 11807031

14. S. Cordes, C.A. Frank, G. Garriga, The C. elegans MELK ortholog PIG-1 regulates cell size asymmetry and daughter cell fate in asymmetric neuroblast divisions. Development 133, 2747–2756 (2006). doi: 10.1242/dev.02447; pmid: 16774992

15. L. Chen et al., Establishment of signaling interactions with cellular resolution for every cell cycle of embryogenesis. Genetics 209, 37–49 (2018). doi: 10.1534/genetics.118.300820; pmid: 29567658

16. J. Cao, M.-K. Wong, Z. Zhao, H. Yan, 3DMMS: Robust 3D membrane morphological segmentation of C. elegans embryo. BMC Bioinform. 20, 176 (2019). doi: 10.1186/s12859-019-2720-x; pmid: 30961566

17. R. Xiong, K. Sugioka, Improved 3D cellular morphometry of Caenorhabditis elegans embryos using a refractive index matching medium. PLoS ONE 15, e0238955 (2020). doi: 10.1371/journal.pone.0238955; pmid: 32997668

18. W. Thiels, B. Smeets, M. Cuvelier, F. Caroti, R. Jelier, spheresDT/Mpacts-PiCS: Cell tracking and shape retrieval in membrane-labeled embryos. Bioinformatics 37, 4851–4856 (2021). doi: 10.1093/bioinformatics/btab557; pmid: 34329378

19. A.D. Chisholm, J. Hardin, Epidermal morphogenesis. WormBook, 1–22 (2005). doi: 10.1895/wormbook.1.35.1; pmid: 18050408

20. J.I. Murray, Z. Bao, T.J. Boyle, R.H. Waterston, The lineaging of fluorescently-labeled Caenorhabditis elegans embryos with StarryNite and AceTree. Nat. Protoc. 1, 1468–1476 (2006). doi: 10.1038/nprot.2006.222; pmid: 17406437

21. X. Li et al., Systems properties and spatiotemporal regulation of cell position variability during embryogenesis. Cell Rep. 26, 313–321 (2019). doi: 10.1016/j.celrep.2018.12.052; pmid: 30625313

22. V.W.S. Ho et al., Systems-level quantification of division timing reveals a common genetic architecture controlling asynchrony and fate asymmetry. Mol. Syst. Biol. 11, 814 (2015). doi: 10.15252/msb.20145857; pmid: 26063786

23. R. Jelier, A. Kruger, J. Swoger, T. Zimmermann, B. Lehner, Compensatory cell movements confer robustness to mechanical deformation during embryonic development. Cell Syst. 3, 160–171 (2016). doi: 10.1016/j.cels.2016.07.005; pmid: 27524104

24. R. Fickentscher, M. Weiss, Physical determinants of asymmetric cell divisions in the early development of Caenorhabditis elegans. Sci. Rep. 7, 9369 (2017). doi: 10.1038/s41598-017-09690-4; pmid: 28839200

25. A. Wang et al., A novel deep learning-based 3D cell segmentation framework for future image-based disease detection. Sci. Rep. 12, 342 (2022). doi: 10.1038/s41598-021-04048-3; pmid: 35013443

26. D. Eschweiler, R.S. Smith, J. Stegmaier, Robust 3d cell segmentation: Extending the view of Cellpose. 2022 IEEE International Conference on Image Processing (ICIP), 191–195 (2022). doi: 10.1109/ICIP46576.2022.9897942

27. M. Weigert, U. Schmidt, R. Haase, K. Sugawara, G. Myers, Star-con27aseline27ledra for 3D object detection and segmentation in microscopy. Proceedings of the IEEE/CVF Winter Conference on Applications of Computer Vision (WACV), 3655–3662 (2020). doi: 10.1109/WACV45572.2020.9093435

28. F. Milletari, N. Navab, S.-A. Ahmadi, V-Net: Fully convolutional neural networks for volumetric medical image segmentation. 2016 Fourth International Conference on 3D Vision (3DV), 565–571 (2016). doi: 10.1109/3DV.2016.79

29. J.E. Sulston, J.G. White, “Parts list” in “The Nematode Caenorhabditis elegans” edited by W.B. Wood. Cold Spring Harbor Laboratory Press, 415–431 (1988). url: https://www.wormatlas.org/celllistsulston.htm

30. K. Driscoll, G.M. Stanfield, R. Droste, H.R. Horvitz, Presumptive TRP channel CED-11 promotes cell volume decrease and facilitates degradation of apoptotic cells in Caenorhabditis elegans. Proc. Natl. Acad. Sci. U. S. A. 114, 8806–8811 (2017). doi: 10.1073/pnas.1705084114; pmid: 28760991

31. R. Jankele, R. Jelier, P. Gönczy, Physically asymmetric division of the C. elegans zygote ensures invariably successful embryogenesis. eLife 10, e61714 (2021). doi: 10.7554/eLife.61714; pmid: 33620314

32. Z.F. Altun, D.H. Hall, Epithelial system, hypodermis. WormAtlas (2009). doi: 10.3908/wormatlas.1.13

33. C.J. Thorpe, A. Schlesinger, J.C. Carter, B. Bowerman, Wnt signaling polarizes an early C. elegans blastomere to distinguish endoderm from mesoderm. Cell 90, 695–705 (1997). doi: 10.1016/s0092-8674(00)80530-9; pmid: 9288749

34. A. Asan, S.A. Raiders, J.R. Priess, Morphogenesis of the C. elegans intestine involves axon guidance genes. PLoS Genet. 12, e1005950 (2016). doi: 10.1371/journal.pgen.1005950; pmid: 27035721

35. Z.F. Altun, D.H. Hall, Muscle system, somatic muscle. WormAtlas (2009). doi: 10.3908/wormatlas.1.7

36. G. Guan et al., Multilevel regulation of muscle-specific transcription factor hlh-1 during Caenorhabditis elegans embryogenesis. Dev. Genes Evol. 230, 265–278 (2020). doi: 10.1007/s00427-020-00662-9; pmid: 32556563

37. R. Fickentscher, S.W. Krauss, M. Weiss, Anti-correlation of cell volumes and cell-cycle times during the embryogenesis of a simple model organism. New J. Phys. 20, 113001 (2018). doi: 10.1088/1367-2630/aaea91

38. A. Sethi et al., A caspase-RhoGEF axis contributes to the cell size threshold for apoptotic death in developing Caenorhabditis elegans. PLoS Biol. 20, e3001786 (2022). doi: 10.1371/journal.pbio.3001786; pmid: 36201522

39. F.K. Nelson, D.L. Riddle, Functional study of the Caenorhabditis elegans secretory-excretory system using laser microsurgery. J. Exp. Zool. 231, 45–56 (1984). doi: 10.1002/jez.1402310107; pmid: 6470649

40. M. Buechner, D.H. Hall, H. Bhatt, E.M. Hedgecock, Cystic canal mutants in Caenorhabditis elegans are defective in the apical membrane domain of the renal (excretory) cell. Dev. Biol. 214, 227–241 (1999). doi: 10.1006/dbio.1999.9398; pmid: 10491271

41. Z. Zhao, L. Fang, N. Chen, R.C. Johnsen, L. Stein, D.L. Baillie, Distinct regulatory elements mediate similar expression patterns in the excretory cell of Caenorhabditis elegans. J. Biol. Chem. 280, 38787–38794 (2005). doi: 10.1074/jbc.M505701200; pmid: 16159881

42. Z.F. Altun, D.H. Hall, Excretory system. WormAtlas (2009). doi: 10.3908/wormatlas.1.17

43. H.M. Ellis, H.R. Horvitz, Genetic control of programmed cell death in the nematode C. elegans. Cell 44, 817–829 (1986). doi: 10.1016/0092-8674(86)90004-8; pmid: 3955651

44. D.M. Eisenmann, Wnt signaling. WormBook (2005). doi: 10.1895/wormbook.1.7.1; pmid: 18050402

45. A.L. Zacharias, T. Walton, E. Preston, J.I. Murray, Quantitative differences in nuclear β-catenin and TCF pattern embryonic cells in C. elegans. PLoS Genet. 11, e1005585 (2015). doi: 10.1371/journal.pgen.1005585; pmid: 26488501

46. P. A. Yushkevich, Y. Gao, G. Gerig, ITK-SNAP: An interactive tool for semi-automatic segmentation of multi-modality biomedical images. 2016 38th Annual International Conference of the IEEE Engineering in Medicine and Biology Society (EMBC), 3342–3345 (2016). doi: 10.1109/EMBC.2016.7591443

47. A. Ajduk, M. Zernicka-Goetz, Polarity and cell division orientation in the cleavage embryo: From worm to human. Mol. Hum. Reprod. 22, 691–703 (2016). doi: 10.1093/molehr/gav068; pmid: 26660321

48. M. Cuvelier, J. Vangheel, W. Thiels, H. Ramon, R. Jelier, B. Smeets, Stability of asymmetric cell division: A deformable cell model of cytokinesis applied to C. elegans. Biophys. J. 122, 1858–1867 (2023). doi: 10.1016/j.bpj.2023.04.017.

49. G. Guan, M.-K. Wong, Z. Zhao, L.-H. Tang, C. Tang, Volume segregation programming in a nema’ ode’ s early embryogenesis. Phys. Rev. E 104, 054409 (2021). doi: 10.1103/PhysRevE.104.054409; pmid: 34942757

50. X. Kuang et al., Computable early Caenorhabditis elegans embryo with a phase field model. PLoS Comput. Biol. 18, e1009755 (2022). doi: 10.1371/journal.pcbi.1009755; pmid: 35030161

## (v) REFERENCES AND NOTES

1. J. Schindelin et al., Fiji: An open-source platform for biological-image analysis. Nat. Methods 9, 676–682 (2012). doi: 10.1038/nmeth.2019; pmid: 22743772

2. Blender Online Community, Blender - A 3D modelling and rendering package. Blender Foundation (2023). url: https://www.blender.org

3. L. Chen et al., Establishment of signaling interactions with cellular resolution for every cell cycle of embryogenesis. Genetics 209, 37–49 (2018). doi: 10.1534/genetics.118.300820; pmid: 29567658

4. Z. Zhao, L. Fang, N. Chen, R.C. Johnsen, L. Stein, D.L. Baillie, Distinct regulatory elements mediate similar expression patterns in the excretory cell of Caenorhabditis elegans. J Biol. Chem. 280, 38787–38794 (2005). doi: 10.1074/jbc.M505701200; pmid: 16159881

5. J. Cao et al., Establishment of a morphological atlas of the Caenorhabditis elegans embryo using deeplearning-based 4D segmentation. Nat. Commun. 11, 6254 (2020). doi: 10.1038/s41467-020-19863-x; pmid: 33288755

6. M.D. Abramoff, P.J. Magalhaes, S.J. Ram, Image processing with ImageJ. Biophotonics Int 11, 36–42 (2004). url: https://imagej.nih.gov/ij/docs/pdfs/Image_Processing_with_ImageJ.pdf

7. J.I. Murray, Z. Bao, T.J. Boyle, R.H. Waterston, The lineaging of fluorescently-labeled Caenorhabditis elegans embryos with StarryNite and AceTree. Nat. Protoc. 1, 1468–1476 (2006). doi: 10.1038/nprot.2006.222; pmid: 17406437

8. Z. Bao, J.I. Murray, T. Boyle, S.L. Ooi, M.J. Sandel, R.H. Waterston, Automated cell lineage tracing in Caenorhabditis elegans. Proc. Natl. Acad. Sci. U. S. A. 103, 2707–2712 (2006). doi: 10.1073/pnas.0511111103; pmid: 16477039

9. B. Katzman, D. Tang, A. Santella, Z. Bao, AceTree: A major update and case study in the long term maintenance of open-source scientific software. BMC Bioinform. 19, 121 (2018). doi: 10.1186/s12859-018-2127-0; pmid: 29618316

10. E. Still, P. Hosemann, Alpha shape analysis: Extracting composition, surface area, and volume post clustering, Microsc. Microanal. 26, 2080–2082 (2020). doi: 10.1017/S1431927620020371

11. O. Ronneberger, P. Fischer, T. Brox, U-Net: Convolutional networks for biomedical image segmentation. Medical Image Computing and Computer-Assisted Intervention (MICCAI) 9351, 234–241 (2015). doi: 10.1007/978-3-319-24574-4_28

12. C. Chen, X. Liu, M. Ding, J. Zheng, J. Li, 3D dilated multi-fiber network for real-time brain tumor segmentation in MRI. Medical Image Computing and Computer Assisted Intervention (MICCAI) 11766, 184–192 (2019). doi: 10.1007/978-3-030-32248-9_21

13. S. van der Walt et al., scikit-image: Image processing in Python. PeerJ 2, e453 (2014). doi: 10.7717/peerj.453; pmid: 25024921

14. M.A. Khan et al., Brain tumor detection and classification: A framework of marker-based watershed algorithm and multilevel priority features selection. Microsc. Res. Tech. 82, 909–922 (2019). doi: 10.1002/jemt.23238; pmid: 30801840

15. A. Wang et al., A novel deep learning-based 3D cell segmentation framework for future image-based disease detection. Sci. Rep. 12, 342 (2022). doi: 10.1038/s41598-021-04048-3; pmid: 35013443

16. D. Eschweiler, R.S. Smith, J. Stegmaier, Robust 3d cell segmentation: Extending the view of Cellpose. 2022 IEEE International Conference on Image Processing (ICIP), 191–195 (2022). doi: 10.1109/ICIP46576.2022.9897942

17. M. Weigert, U. Schmidt, R. Haase, K. Sugawara, G. Myers, Star-convex polyhedra for 3D object detection and segmentation in microscopy. Proceedings of the IEEE/CVF Winter Conference on Applications of Computer Vision (WACV), 3655–3662 (2020). doi: 10.1109/WACV45572.2020.9093435

18. F. Milletari, N. Navab, S.-A. Ahmadi, V-Net: Fully convolutional neural networks for volumetric medical image segmentation. 2016 Fourth International Conference on 3D Vision (3DV), 565–571 (2016). doi: 10.1109/3DV.2016.79

19. B. Sönnichsen et al., Full-genome RNAi profiling of early embryogenesis in Caenorhabditis elegans. Nature 434, 462–469 (2005). doi: 10.1038/nature03353; pmid: 15791247

20. R.A. Green et al., A high-resolution C. elegans essential gene network based on phenotypic profiling of a complex tissue. Cell 145, 470–482 (2011). doi: 10.1016/j.cell.2011.03.037; pmid: 21529718

21. P. A. Yushkevich, Y. Gao, G. Gerig, ITK-SNAP: An interactive tool for semi-automatic segmentation of multi-modality biomedical images. 2016 38th Annual International Conference of the IEEE Engineering in Medicine and Biology Society (EMBC), 3342–3345 (2016). doi: 10.1109/EMBC.2016.7591443

22. J.E. Sulston, E. Schierenberg, J.G. White, J.N. Thomson, The embryonic cell lineage of the nematode Caenorhabditis elegans. Dev. Biol. 100, 64–119 (1983). doi: 10.1016/0012-1606(83)90201-4; pmid: 6684600

23. X. Li et al., Systems properties and spatiotemporal regulation of cell position variability during embryogenesis. Cell Rep. 26, 313–321 (2019). doi: 10.1016/j.celrep.2018.12.052; pmid: 30625313

